# Time-scaled phylogenetic analysis of extant Lamiinae (Coleoptera, Cerambycidae) species of East of Marmara Basin, Türkiye, and their evolutionary similarity with the Eurasian congeners

**DOI:** 10.1101/2022.09.10.507409

**Authors:** Havva Kübra Soydabaş-Ayoub, Fevzi Uçkan

## Abstract

The subfamily Lamiinae (Cerambycidae, Coleoptera) is striking due to its morphological diversity and species richness with intricate phylogenetic relationships. We inferred the phylogeny and evolutionary history of extant species of East of Marmara Basin, Türkiye, from the tribes Acanthocinini, Acanthoderini, Agapanthiini, Batocerini, Dorcadionini, Lamiini, Mesosini, Monochamini, Phytoeciini, Phrynetini, Pogonocherini (including Exocentrini) and Saperdini using partial mitochondrial cytochrome *c* oxidase-I (*COI*) and *16S rRNA* and nuclear *28S rRNA* gene regions (2257 base pair alignment length). The most recent common ancestor (MRCA) of Lamiinae members included in the analyses was dated ∼127 million years ago (Mya) in the Cretaceous. The MRCA of Dorcadionini, Lamiini and Monochamini was younger than the common ancestors of the other close tribes. There was a concurrence between resolutions of Maximum Likelihood (ML) and Bayesian analyses on the affiliations of Dorcadionini and Monochamini to Lamiini and the proximity of Batocerini to Lamiini, Acanthocinini to Acanthoderini, Phrynetini to Pogonocherini, and Phytoeciini to Saperdini. The *COI*-based Neighbor-Joining and ML gene trees suggest that the closest relatives of the extant Lamiinae species of East of Marmara Basin were the European conspecifics or congeners. Moreover, *Paraleprodera* and *Lamia* (Lamiini) were sisters to *Imantocera* (Gnomini), *Oberea* (Obereini) to *Phytoecia* Phytoeciini), and *Hippopsis* (Agapanthiini) to *Omosarotes singularis* Pascoe, 1860 (Acanthomerosternoplini). Our results support Dorcadionini, Gnomini and Monochamini as synonyms of Lamiini; and Obereini and Phytoeciini of Saperdini and suggest that the emergence of the living tribes included in this study was during Paleogene, and their intrageneric diversifications occurred during Cenozoic, mostly Neogene.

## 1. Introduction

Cerambycidae, also known as long-horned wood-boring beetles, is the species-rich family of the superfamily Chrysomeloidea (Coleoptera, Phytophaga), with about 34,000 described species; among them, around 20,000 extant species belong to subfamily Lamiinae (Tavakilian and Chevillotte, 2022). The family comprises mainly herbivorous beetles distributed in all biogeographic realms except Antarctica (Monné et al., 2017). Some species are considered pests of wood due to their larval feeding habits; however, beyond this, they have pivotal roles in sustaining the harmony of the forest ecosystem, such as pollination and decaying wood (Linsley, 1959).

The prominent dependence of cerambycids on the plant has led to investigations of their evolution as a part of the herbivorous beetles (Phytophaga) clade, whose over-diversification has been mainly ascribed to their adaptive radiation (Farrell, 1998; McKenna et al.; 2015; Zhang et al., 2018; McKenna et al. 2019). According to Farrell (1998), basal lineages of cerambycids emerged in the Jurassic, and the ancestral lineages were associated with gymnosperms. The enhanced net diversification rate of primitively angiosperm-associated taxa is related to a series of adaptive radiations of beetles onto angiosperm.

In contrast to him, Gómez-Zurita et al. (2007) suggested that the origin of the Chrysomelidae is 73–79 Mya and the origin of cerambycids is Late Cretaceous about 80 Mya, thus cannot be associated with diversification of Early Cretaceous angiosperm lineage. Hunt et al. (2017) placed the origin of Coleoptera at 285 Mya, dated the origin of cerambycids at 203 Mya, and dated the first appearance of most of the modern lineages of beetles in the Jurassic. Despite similar dating with Farrell (1998), Hunt et al. (2017) concluded that the underlying reason for the success of beetles is neither the net diversification rate nor the Cretaceous rise of angiosperms, instead sustained diversification in various niches. Then the divergence time of Chrysomeloidea was recalibrated by Wang et al. (2014) due to a discovery of the known earliest cerambycid fossil, *Cretoprionus liutiaogouensis* Wang, et al., 2013 (Prioninae) from the Early Cretaceous of China. According to their estimation, the origin of modern Cerambycidae was the Late Triassic (∼210 Mya), older than the estimates of Gómez-Zurita et al. (2007), and all subfamilies arose by the mid-Cretaceous. Then, again, the estimation of the emergence time of the family was shifted to the beginning of the Early Cretaceous around 145 Mya by Zhang et al. (2018). McKenna et al. (2019) dated the origin of Coleoptera to the Carboniferous about 327 Mya. They also suggested that the diversification rate within Polyphaga increased in the Late Carboniferous by about 305 Mya. The origins of most of the extant beetle families, including Cerambycidae, appeared before the end of the Cretaceous, and the emergence of cerambycids was around 160 Mya (McKenna et al., 2019). Nie et al. (2021) dated the emergence of the ancestor of the crown group of the subfamily Lamiinae at 132.0 Mya.

Besides evolutionary relationships, the studies inferring phylogenetic relationships of the subfamilies of Cerambycidae are primarily focused on the higher taxa. The studies rely on either morphological or molecular synapomorphic characters or both (Napp, 1994; Švácha and Lawrence, 2014; Wei et al., 2014; Haddad et al., 2018; Nie et al., 2021), have reached almost complete unanimity on the monophyly of the subfamily Lamiinae. However, the morphological and molecular studies conflict in who is the sister clade of this subfamily. In contrast to the previous morphological studies (Villiers, 1978; Napp, 1994) that suggest Cerambycinae as the sister of Lamiinae, molecular phylogenetic analysis (Gómez-Zurita et al., 2007; Marvaldi et al., 2009; Haddad et al., 2018; Nie et al., 2021) have concurrence that Spondylidinae is the sister clade of Lamiinae.

Recently, some studies took initial steps to resolve the tribal-level relationships of Lamiinae based on genetic data. de Santana Souza et al. (2020) inferred using mitochondrial and nuclear marker DNA sequences that among the phylogenetically analyzed 46 tribes, Astathini, Batocerini, Ceroplesini, Colobotheini, Compsosomatini, Dorcadionini, Lamiini, Mesosini, Obereini, and Polyrhaphidini are monophyletic; but, some of widely distributed or diverse tribes such as Acanthocinini, Acanthoderini, Agapanthiini, Apomecynini, Desmiphorini, Dorcaschematini, Enicodini, Hemilophini, Monochamini, Onciderini, Parmenini, Phytoeciini, Pogonocherini, Pteropliini, and Saperdini are polyphyletic. Ren et al. (2021) presented the sisterships of Ceroplesini and Agapanthiini, Mesosini and Pteropliini, Saperdini and Phytoeciini, Batocerini and Lamiini, and Apomecynini and Acanthocinini based on analyzed 13 protein-coding genes of mitogenomes of 12 tribes’ representatives. Ashman et al. (2022) reported that Acanthocinini, Apomecynini, Parmenini, Desmiphorini, Lamiini and Pteropliini are polyphyletic, relying on the phylogenetic analysis of hundreds of nuclear gene regions, including representatives of 14 tribes, mostly Australian. The present study aims to figure out the evolutionary closeness of the Lamiinae species that presently occur in the East of Marmara Basin, Türkiye, with their Eurasian relatives using partial mitochondrial cytochrome *c* oxidase-I (*COI*) gene region and to contribute to phylogenetic relationships and the evolutionary history of tribes Acanthocinini, Acanthoderini, Agapanthiini, Monochamini, Phytoeciini, Pogonocherini, and Saperdini using partial mitochondrial *COI* and *16S rRNA* and nuclear *28S rRNA* gene regions.

## 2. Materials and Methods

Specimens were collected from timber yards, wood processing plants, suburban areas and forests between 2016 and 2019 in East of Marmara Basin, Türkiye (Supplementary Table S1), a remarkable region for insect biodiversity due to its geographic position and climatic (Çakmak et al., 2019); vegetative (Atak et al., 2021), and mercantile (Soydabaş-Ayoub et al., 2022) characteristics. Ethanol and α-pinene tubes were used within three funnel traps. Insect nets were used for flower-visiting species. *Phryneta leprosa* (Castilla-borer) was intercepted in Derince port on timber imported from Cameroon. *Batocera rufomaculata* (tropical fig-borer) was caught coincidentally in Diyarbakır, Türkiye. Morphological identifications were carried out with the guidance of Breuning (1951), Bílý and Mehl (1989), Bense (1995), Sama (2002), Wallin, Nylander, and Kvamme (2009), and Özdikmen (2013) under a stereomicroscope (Olympus SZ51, Japan) enhanced by Olympus 110AL2X WD38 auxiliary macro lens. Specimens were stored in 99% ethanol at -20 °C pending DNA extraction.

Depending on the specimen sizes, muscle tissues of coxa or antennae were shredded on disposable sterile slides by disposable scalpels. Then transferred into a microcentrifuge tube including the lysis buffer (8 mM dithiothreitol (DTT), 2% sodium dodecyl sulphate (SDS), 100 mM NaCl, 3 mM CaCl2, dissolved in 100 mM Tris buffer (pH 8) and 200 µg/mL proteinase K) (Soydabaş-Ayoub, 2021) and kept in Eppendorf, Germany, Thermomixer 5350 heater at 37°C until whole tissue whittle down. Genomic DNA was extracted by following the phenol:chloroform:isoamyl alcohol (25:24:1) protocol (Sambrook and Russell, 2006).

Polymerase chain reaction (PCR) products were amplified in a total volume of 25µL with QuickLoad® Taq 2X Master Mix (New England Biolabs Inc., USA, catalogue number M0271L) and 0.2 µM primer for each pair, with the addition of 0.06 mg/mL bovine serum albumin. The primer pairs HCO2198–LCO1490 (Folmer et al., 1994) for the mitochondrial *COI* gene region, LR-J-12887-LR-N-13398 (Yoon et al., 2001) for mitochondrial *16S rRNA* gene region, and LSU D1, D2 fw1: 56-74 and LSU D1, D2r rev2: 1048-1067 (Sonnenberg et al., 2007) for nuclear *28S rRNA* gene regions were used in amplification. Mitochondrial *COI* and *16S rRNA* gene regions were amplified by using the same thermocycling parameters as 1 min at 95 °C, five cycles of 30 s at 95 °C, 1 min at 46 °C, 1 min at 72 °C, 30 cycles of 30 s at 95 °C, 1 min at 51°C, 1 min at 72 and 10 min at 72 °C (Çakmak et al., 2020). The thermocycling parameters used for amplification of the nuclear *28S rRNA* gene region were 1 min at 95 °C, four cycles of 30 s at 95 °C, 1 min at 57,3 °C, 1 min at 72 °C, and 20 cycles of 30 s at 94 °C, 1 min at 61,7 °C, 1 min at 72 °C and 10 min at 72 °C (Soydabaş-Ayoub et al., 2022). ExoSAP-IT™ Clean-Up Reagent was used for purification according to manufacturer instructions. Sequencing was performed for both directions by ABI 3730XL DNA Analyzer (Applied Biosystems, Foster City, CA) at Macrogen Laboratory, Holland.

Geneious Prime v2019.2.1 (Kearse et al., 2012) was used for processing raw data. Prior to BLAST searches of the contigs of high-quality (HQ% >90) chromatograms, the primer regions and the leftover extensions following primer sequences were trimmed.

The heterozygote bases in nuclear gene sequences were named based on the IUPAC (International Union of Pure and Applied Chemistry) base nomenclature code system. A total of 143 obtained sequences were deposited in GenBank under the accession numbers OP279135-OP279183, OP279535-OP279581 and OP279486-OP279532 for mitochondrial *COI*, *16S rRNA* and nuclear *28S rRNA* gene regions, respectively (Supplementary Table S1). Two datasets were prepared for the phylogenetic analysis. The first one was a global mitochondrial *COI* sequence dataset of the subfamily Lamiinae, which was configured by combining the sequences obtained in this study and retrieved from the BOLD taxonomy archive. To improve the dataset’s reliability, sequences shorter than 658 bp were discarded. A representative was selected for each available species from each zoogeographic region based on Löbl and Smetana (2010) (i.e., Neotropical, Nearctic, Afrotropical, Oriental Australian, Palearctic: Europe, Asia, North Africa) (Supplementary Table S2).

The second dataset consisted of two mitochondrial and one nuclear marker sequences; all were produced in this study (Supplementary Table S1). The mitochondrial *COI*, *16S rRNA*, and nuclear *28S rRNA* gene regions were concatenated in Geneious Prime v2019.2.1 (Kearse et al., 2012) and aligned by MUSCLE v3.8.425 (Edgar, 2004). Model selections were carried out by PartitionFinder 2 (Lanfear et al., 2017), the basis of AICc, by greedy search.

Phylogenetic trees were obtained using Maximum Likelihood (ML) and Neighbor-Joining (NJ) approaches. ML analysis was conducted by using the PhyML (Guindon et al., 2010) plug-in of Geneious Prime v2021 for spliced dataset (Supplementary Table S1) of three gene regions (2257 bp) with 34 terminal, produced in this study; and a global dataset of the mitochondrial *COI* gene region, including the sequences produced in this study (Supplementary Table S1) and retrieved from databases (658 bp) with a total of 441 terminal (Supplementary Table S2). For both datasets, ML analyses were carried out by 10,000 bootstrap pseudo-replicates with estimated parameters and the Tamura-Nei (TN93) substitution model for unpartitioned data. The outgroups used in phylogenetic analyses were selected from the closest relatives according to Haddad et al. (2018): *Spondylis buprestoides* (Linnaeus, 1758) (Spondylidinae: Spondylidini), *Arhopalus rusticus* (Linnaeus, 1758) (Spondylidinae: Asemini). NJ analysis was conducted by tree builder in Geneious Prime v2021 with 10,000 bootstrap pseudo-replicates and TN93 substitution model for only the global *COI* dataset (Supplementary Table S1).

The time-scaled Bayesian analysis was conducted by BEAST v1.10.4 (Drummond and Rambaut, 2007) under the Yule speciation process (Gernhard, 2008) and uncorrelated relaxed clock model with lognormal distribution (Drummond et al., 2006). Molecular clock estimation was calibrated considering fossil records and chronograms of recent studies (McKenna et al., 2019; Nie et al., 2021; Ashman et al., 2022). The normal distribution with ±5% standard deviation was used for calibrating the root ages. The mean ages of the most common recent ancestor (MRCA) were set to 147.03, 138.1, and 132.0 Mya for the lamiine clade (Lamiinae+Spondylidinae), Spondylidinae and Laminae, respectively based on Nie et al. (2021). The root age (crown lamiine clade) was constrained to 180 Mya (McKenna et al., 2019). The lower bounds of MRCAs of Spondylidinae and Laminae were truncated to 34 Mya and 57 Mya, respectively, relying on fossil records of *Arhopalus pavitus* Cockerell 1927 from Eocene of Colorado (Cockerell, 1927) and *Palaeoncoderes* and *Prolamioides* from Paleocene of France (Piton and Théobald, 1937).

The Jukes-Cantor (JC) substitution model was selected after trial runs to determine the best-fitting model since it provided higher posterior probabilities than other substitution models available in BEAST v1.10.4 (HKY, GTR, and TN93). Auto-optimized classical operator mix parameters were used for two Bayesian analyses, run for 100 million MCMC generations, and sampled every 1000 steps; of these, 10% were discarded as burn-in. LogCombiner v1.10.4 merged the outputs, and the maximum clade credibility (MCC) trees resulting from each run were interpreted by TreeAnnotator v1.10.4 in the BEAST package. Tracer 1.7 (Rambaut et al., 2018) was used to assess if the implemented parameters provide effective sample sizes (ESSs). The MCC trees were annotated in FigTree v1.4.4 (Rambaut, 2014), and the visual adjustments were performed in CorelDraw v.1512.

## 3. Results

### 3.1. Phylogenetic Analysis

The hypothetical phylograms resulting from the NJ and ML analyses of the global dataset of *COI* barcode region adumbrated relationships, particularly at the generic level. All specimens obtained in this study from the East of Marmara Basin were clustered with the European conspecifics, congeners or relatives (Supplementary Figure S1, Supplementary Figure S2). The phylogram hypothesized by the ML approach of spliced datasets of mitochondrial *COI* and *16S rRNA* and nuclear *28S rRNA* gene regions (Supplementary Table S1) of Lamiinae schemed a pattern as follows: The first main branch including [(Lamiini+Dorcadionini) + (Monochamiini+Batocerini)] + Mesosini, the second Phrynetini+Pogonocherini (including Exocentrini), the third Agapanthiini, fourth Acanthocinini+Acanthoderini, and the last one Saperdini+Phytoeciini (Figure 1). For traceability, we mostly used generic names of the representatives of tribes in the following text.

**Figure 1.**
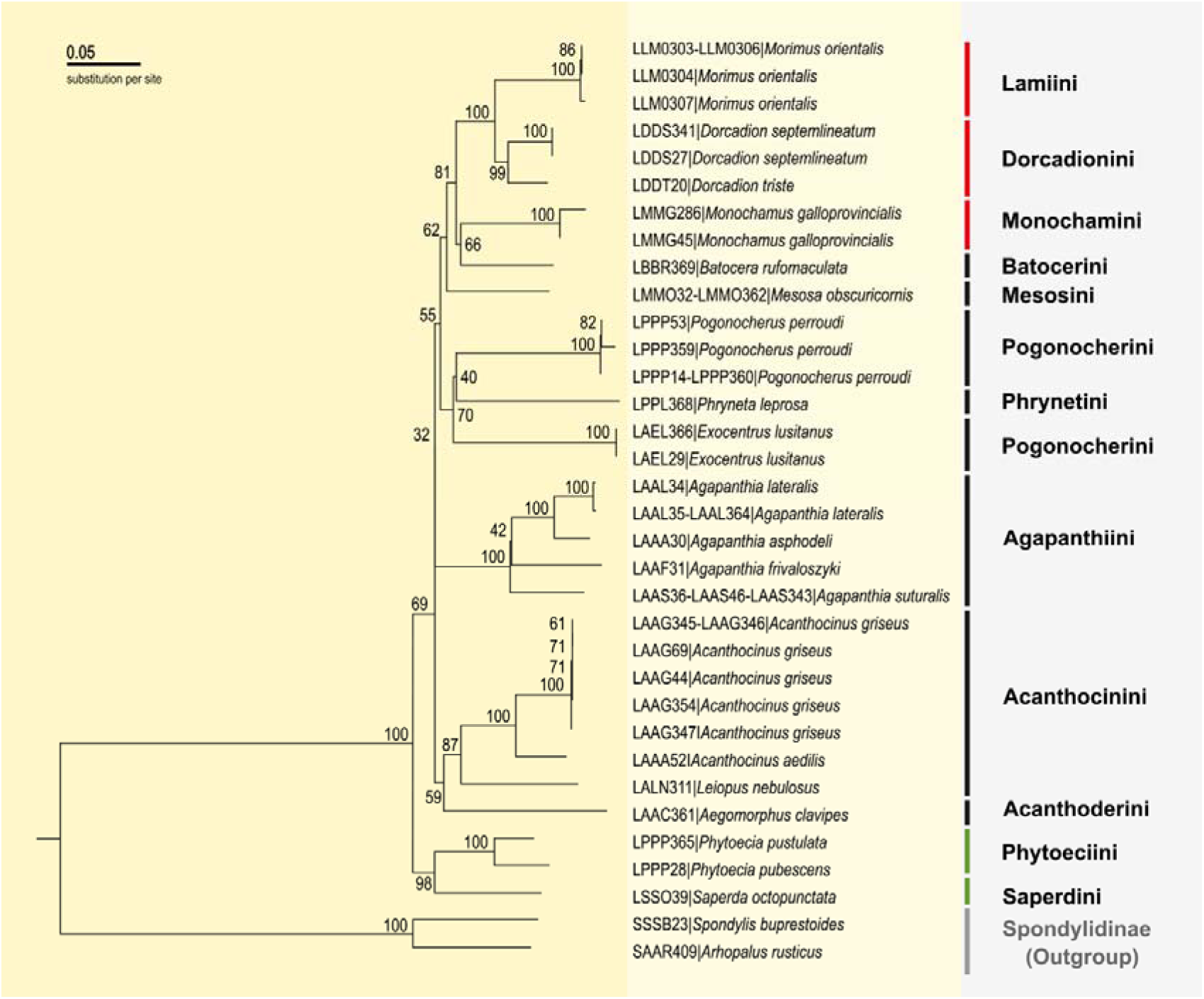
Phylogenetic relationships recovered by maximum likelihood (ML) analysis of spliced datasets of mitochondrial *COI* and *16S rRNA* and nuclear *28S rRNA* gene regions. Bootstrap supports are indicated beside nodes.

The *COI*-based NJ and ML gene trees supported the affiliation of the genera *Morimus* (Lamiini) and *Dorcadion* (Dorcadionini), besides the genus *Iberodorcadion* (Dorcadionini) nested in the same group, sister to *Morimus+Dorcadion* branch. The other genera splinted from the same node in the NJ tree were *Paraleprodera* and *Lamia* (Lamiini), *Imantocera* (Gnomini), *Batocera* and *Apriona* (Batocerini) (Supplementary Figure S1 and Supplementary Figure S2). *Peblephaeus* (Lamiini) was stated at the basal-most of the group at the NJ tree (Supplementary Figure S1). *Mesosa obscuricornis* was clustered with congeners and *Synaphaeta* (Mesosini) and *Acalolepta* (Lamiini) (Supplementary Figure S1 and Supplementary Figure S2).

The Phrynetini+Pogonocherini branch was recovered as the sister group of Lamiini and the relatives mentioned above (Figure 1). The genus *Exocentrus*, which is a member of the tribe Pogonocheriini (or Exocentrini), was nested within this cluster with a 70% bootstrap support at the ML tree of the spliced datasets of mitochondrial *COI* and *16S rRNA* and nuclear *28S rRNA* gene regions (Figure 1).

At the *COI*-based NJ and ML trees, the members of the genus *Pogonocherus* were clustered all together, split away from other members of Pogonocherini. This cluster was the basal-most of the *COI*-based ML tree. The two genera in Pogonocherini, *Ecyrus* and *Exocentrus*, were clustered with *Rosalba sp*. (Apomecynini) and *Bactriola* sp. (Forsteriini) at the *COI*-based ML tree Supplementary Figure S2). The genus *Phryneta* was clustered with the members of the genera *Anaesthetis* and *Pseudanaesthetis* from the tribe Desmiphorini (Supplementary Figure S2), while *Exocentrus* was clustered with *Desmiphora* (Desmiphorini) at the *COI*-based NJ tree (Supplementary Figure S1).

The tribe Agapanthiini was represented by the genus *Agapantia* at the ML tree of the spliced datasets of mitochondrial *COI* and *16S rRNA* and nuclear *28S rRNA* gene regions, stated at the base of the groups mentioned above with trivial statistical support (Figure 1). At the *COI*-based NJ and *COI*-based ML trees, two other species from Agapanthiini, *Calamobius filum* (Rossi, 1790) and *Pothyne virginalis* Takakuwa and Kusama, 1979 were clustered in the *Agapantia* genus group (Supplementary Figure S1, Supplementary Figure S2).

The sole representative of the tribe Acanthoderini *Aegomorphus clavipes*, stated at the base of the Acanthocinini and Acanthoderini group at the ML tree of spliced datasets of mitochondrial *COI* and *16S rRNA* and nuclear *28S rRNA* gene regions (Figure 1). At the *COI*-based NJ and *COI*-based ML trees *Aegomorphus modestus* (Blais, 1817) was joined to *A. clavipes*. They were clustered with other Acanthoderini members, such as genera *Acanthoderes, Paradiscopus, Psapharochrus* and *Steirastoma* (Supplementary Figure S1, Supplementary Figure S2).

The basal-most branch of the ML tree of the spliced datasets of mitochondrial *COI* and *16S rRNA* and nuclear *28S rRNA* gene regions included Saperdini and Phytoeciini (Figure1). At the tree resulting from *COI*-based NJ and *COI*-based ML analysis, the genus *Saperda* group included the members of genera *Eutetrapha*, *Glenea*, *Thyestilla* and *Stenostola* from the tribe Saperdini. In the genus group of *Phytoecia*, the genus *Oberea* (Obereini) was nested. These two groups were stated as sister clades (Supplementary Figure S1, Supplementary Figure S2).

### 3.2. Divergence Time Estimation

The time-scaled Bayesian analysis of the spliced datasets of mitochondrial *COI* and *16S rRNA* and nuclear *28S rRNA* gene regions (Supplementary Table S1) suggested a mostly congruent topology with the ML analysis (Figure 1, Figure 2). The MRCA of Lamiinae rely on the taxa involved in the analysis, raised around the Early Cretaceous, 127.49 Mya (95% highest posterior density (HPD) interval 117.95-137.57 Mya). The common ancestor of the Lamiini and its close relatives (Dorcadionini, Monochamiini, Batocerini and Mesosini appeared 66.77 Mya (95% HPD: 42.34-92.92 Mya) around the Late Cretaceous to Eocene epoch of the Paleogene. Also, appearing times of the earliest common ancestors of the other groups were from Late Cretaceous to the different epochs of the Paleogene. Phrynetini+Pogonocherini (including Exocentrini) and Acanthocinini+Acanthoderini groups emerged in Late Cretaceous to Paleogene 58.85 Mya (HPD: 29.25-87.85 Mya) and 77.71 Mya (95% HPD: 40.68-111.73 Mya), respectively. The MRCA of Saperdini+Phytoeciini emerged in 65.24 Mya (95% HPD: 23.06-111.61 Mya), somewhere from the mid-Cretaceous to the end of Oligocene. While the emerging time of the MRCA of the *Agapanthia* genus group was 54.74 Mya (95% HPD: 27.5-87.9 Mya), the emerging time of the MRCA of the tribes Dorcadionini, Lamiini, and Monochamini was 41.28 Mya (95% HPD: 21.51-63.81 Mya). Besides, the MRCA of these three tribes was younger than all MRCAs of the other tribes (Figure 2). The intrageneric diversification relies on the taxa involved in the analysis that occurred during Cenozoic. The members of *Agapanthia* arose during Paleogene 54.74 Mya (95% HPD: 27.5-87.9 Mya), earlier than *Phytoecia* members 23.13 Mya (95% HPD: 3.91-54.16 Mya.) The last species emerging within our sampling corresponded to the Neogene and occurred within the genus *Dorcadion* 14.34 Mya (95% HPD: 4.04-29.83 Mya).

**Figure 2.**
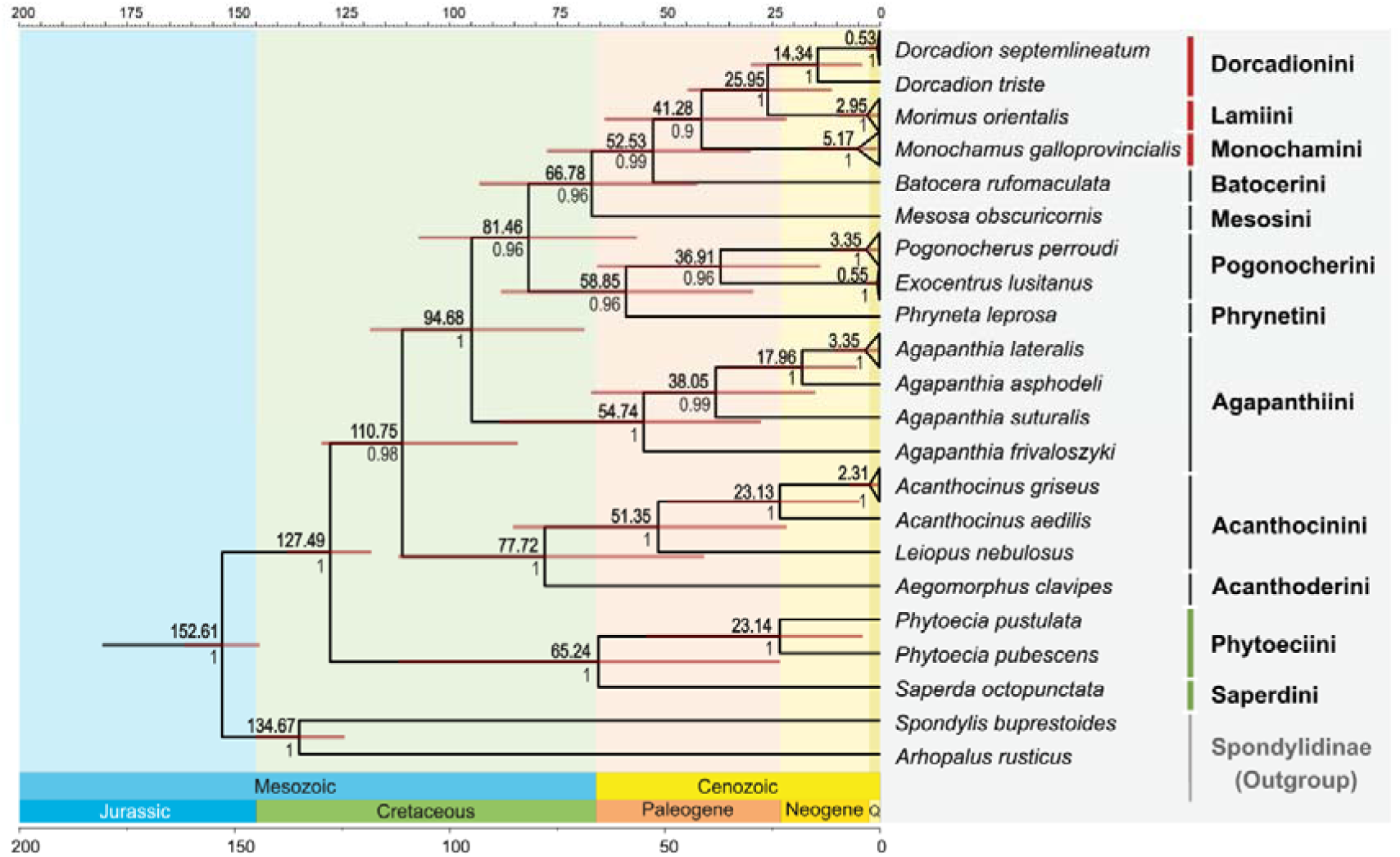
Divergence times recovered by Bayesian analysis of spliced datasets of mitochondrial COI and *16S rRNA* and nuclear *28S rRNA* gene regions. The brown bars correspond to the 95% highest posterior density (HPD) interval; the mean ages are above, and the posterior probabilities are below each node; the scale bars indicate the time in million years; Q: Quaternary.

## 4. Discussion

The previous studies contributing knowledge of the evolutionary history and phylogenetic relationships of Lamiinae are mainly dealing with higher-level relationships and covering only a few laminin taxa (Farrell, 1998; Gómez-Zurita et al., 2007; Hunt et al., 2007; Wang et al., 2014; Haddad et al., 2018; Zhang et al., 2018). Since the inception of diversification of herbivorous beetles has been set on relatively solid ground, the studies are bending to lower taxonomic levels. Phylogeny of Lamiinae, especially at tribal and generic levels, has remarkable progress owing to de Santana Souza et al. (2020), Ashman et al. (2022) and Dascălu et al. (2022). However, the relationships at tribal and generic levels still require intense effort due to the species richness, remarkable diversity and the worldwide distribution of the subfamily. The present study contributes to understanding tribal and generic level phylogenetic relationships and the evolutionary history of Lamiinae using Eurasian species as the sampling allows.

### 4.1. Phylogenetic Relationships

The topologies suggested by the multi-locus ML and the time-scaled BI analyses showed consistency in the clustering of the genera *Morimus* (Phrissomini syn. of Lamiini), *Dorcadion* (Dorcadionini), *Monochamus* (Monochamini) and *Batocera* (Batocerini) (Figure 1, Figure S2). Single-locus *COI*-based NJ and ML trees analyses provided a broader aspect due to covering more taxa, and clustered *Dorcadion* and *Iberodorcadion* (Dorcadionini); *Imantocera* (Gnomini), *Batocera* and *Apriona* (Batocerini) with genera *Lamia*, *Morimus*, *Paraleprodera*, *Blepephaeus* and *Peblephaeus* at (Supplementary Figure S1, Supplementary Figure S2). This propinquity has long been recognized and suggested merging all tribes which have the guise of Monochamini by Breuning (1961) and supported by the following studies (Ohbayashi et al., 2009; Toki and Kubota, 2010; Gorring, 2019; de Santana Souza et al., 2020). Additionally, de Santana Souza et al. (2020) perceived Dorcadionini as the closest relative of Lamiini, despite its apparent difference, which is most probably due to only adaptive characters (Löbl and Smetana, 2010). The close kinship of Lamiini and Dorcadionini was also supported by Giannoulis et al. (2020); however, their results were dissimilar to other studies due to the placement of *Phytoecia* (Phytoeciini) at the base of the Lamiini+Dorcadionini branch together with *Monochamus* (Monochamini). Our results were conformable with the results of de Ashman et al. (2022) and de Santana Souza et al. (2020) on the station of *Monochamus* and *Batocera* closely. Ren et al. (2021) also supported the sisterships of Batocerini and Lamiini. Contemplating the results of the present study and the previous studies mentioned above, the expectations of Lacordaire (1869) and Pascoe (1866) (as per de Santana Souza et al., 2020) regarding the close relationship between Batocerini and Monochamini seem to be plausible, and Dorcadionini, Gnomini, and Monochamini (at least in terms of *Monochamus*) should be revised.

Mesosini split from Lamiini and close relatives mentioned above by the absence of a sharp tubercle or spine at the lateral margin of the pronotum and a simple mesotibia (Bílý and Mehl, 1989; Bense, 1995). However, for both tribes, the first antennal segment has a rounded apex with a carina, which separates them from Saperdini (Bílý and Mehl, 1989). The proximity of Mesosini to the cluster of Lamiini has been shown in the phylogenetic trees of de Santana Souza et al. (2020) and is supported by our results. (Figure 1, Figure 2, Supplementary Figure S1, Supplementary Figure S2). However, unlike de Santana Souza et al. (2020), *Mesosa* was not in the same subcluster as *Saperda* (Saperdini) in our phylogenetic trees. Therefore, the apex structure of the first antennal segment might be evaluated as a mark for the synapomorphy of Mesosini and Lamiini.

Phrynetini, represented by *P. leprosa*, was nested in Pogonocherini in our phylogenetic trees, while it was at the base of the Acanthoderini, according to de Santana Souza et al. (2020)’s results. This tribe is mostly occurring Afrotropic and Indomalaya regions, and its closely related taxa were possibly not included in analyses; thus, it would not be accurate to conclude its relationships.

Our results support the closeness of *Pogonocherus* and *Exocentrus* shown by de Santana Souza et al. (2020) and Ashman et al. (2022). On the other hand, considering our *COI*-based ML tree (Supplementary Figure S2), clusterings of *Ecyrus* and *Exocentrus* of Pogonocherini with *Rosalba* sp. (Apomecynini) and *Bactriola* sp. (Forsteriini), respectively might be a sign of a need for questioning monophyly of Forsteriini by extensive sampling and broader genetic data, in addition to 12 tribes and Apomecynini discussed by de Santana Souza et al. (2020)

The tribes Acanthocinini and Acanthoderini, which share some morphological traits such as antennae lacking long erect hairs and laterally closed joint sockets of middle coxa (Bílý and Mehl, 1989; Bense, 1995), were sister clades in our time-scaled-BI and ML analyses, *A. clavipes*, the sole representative of the Acanthoderini, stated at the base of Acanthocinini, while it was clustered with *P. leprosa* from Phrynetini in de Santana Souza et al. (2020).

Agapanthiini, the 12-segmented tribe of the subfamily, was not monophyletic at the *COI*-based NJ and ML gene trees of the present study; *C. filum* was clustered with the genus *Agapanthia*. In contrast, *Hippopsis* sp. from this tribe was clustered out of this group, with *O. singularis* from the tribe Acanthomerosternoplini (Supplementary Figure S1). Agapanthiini seems polyphyletic at ML tree of de Santana Souza (2020) et al., in terms of *Hippopsis* sp., also.

The closeness of Obereini, Phytoeciini, and Saperdini was shown by de Santana Souza et al. (2020), who used the genera *Oberea* from the tribe Obereini and *Mecas* (*Dylobolus*) and *Phytoecia* from Phytoeciini, and *Glenea, Paraglenea,* and *Saperda* from Saperdini the present study’s *COI*-based NJ and ML gene trees supported their results. The Saperdini group included *Eutetrapha, Glenea, Mecas, Saperda, Stenostola, Thyestilla*, and *Phytoecia* (Supplementary Figure S1, Supplementary Figure S1). Same as Ren et al. (2021), and Ashman et al. (2022), our findings support the suggestion of de Santana Souza et al. (2020) that the tribe Phytoeciini should also be a synonym of Saperdini or all members should be evaluated separately.

### 4.2. Diversification Times

Initial evolutionary history studies rely on genetic data to estimate the emergence of cerambycids from the Late Cretaceous (Gómez-Zurita et al., 2007) to the Early Cretaceous (Wang et al., 2014; Zhang et al., 2018), while recent studies, which were conducted with broader samplings and genetic data, have pointed out earlier emergence time, around late Jurassic to early Cretaceous (Nie et al., 2021; Ashman et al., 2022). Our time-scaled BI analysis dated the crown age ∼153 Mya, at Late Jurassic to Early Cretaceous, compatible with the emergence times presented in previous studies (Farrell, 1998; Wang et al., 2014; Yu et al., 2015; Zhang et al., 2018; McKenna et al., 2019; Nie et al., 2020; Ashman et al., 2022). Ashman et al. (2022) and Nie et al. (2021), whose time estimation studies cover broader laminin samples, have conflicts in some emergence times of MRCA of tribes, probably due to differences between their genomic data, calibration points, and the taxa included in the analyses. Also, our estimations resulted in some discordances due to the same reasons. All three of us are probably missing the most major branching events due to the scarcity of the taxa.

According to the chronogram of Ashman et al. (2022), MRCA of the subfamily Lamiinae emerged at ∼105 Mya; *Acanthocinus griseus* (Acanthocinini) split from the MRCA of Lamiinae and stated at the base of the Lamiinae clade. According to our chronogram, the emergence of the MRCA of the subfamily ∼128 Mya was approximately 50 Mya earlier than the MRCA of Acanthocinini and Acanthoderini, which emerged ∼78 Mya at the late Cretaceous. The split of *A. griseus* and *A. aedilis* was dated ∼23 Mya, earlier than the oldest known fossil species of the genus *Acanthocinus schmidti* Schmidt 1967 3.6 to

2.588 Mya from Pliocene of Germany (Gersdorf, 1976). Other fossil records *Astynomus tertiarius* Kolbe 1888 and *Kallyntrosternidius bucarensis* Vitali, 2009 of Acanthocinini address an earlier date (15.97 to 13.65 Mya) Miocene of Germany and 20.43 to 13.65 Mya Miocene of Dominican Republic (Vitali, 2009), respectively. The fossil records of Acanthoderini, *Acanthoderus lepidus* Heer 1865, *Acanthoderes phrixi* Heer 1847 and *Acanthoderus sepultus* Heer 1865, are known from the Miocene of Croatia and Germany 12.7 to 11.6 Mya (Heer, 1879). However, any study that mentions fossil traces of a common ancestor of Acanthocinini and Acanthoderini has yet to be encountered.

The MRCA of the Lamiini and close relatives, which include *Monochamus* and *Batocera*, appeared at the Eocene epoch of Paleogene, ∼52.53 Mya according to our chronogram, relatively consistent, but a bit earlier than the chronogram of Ashman et al. (2022) and Nie et al. (2021) which point to ∼63 Mya and ∼55 Mya, respectively. This is the youngest clade compared to the other tribes included in this study (Figure 2). The fossil records of Lamiini, *Lamia antiqua* Heer 1879, from Miocene of Germany (12.7 to 11.6 Mya) (Heer, 1879), *Dorcadion bachense* Handschin 1944 from Oligocene of France (28.4 to 23.03 Mya) (Handschin, 1944), and *Dorcadion emeritum* von Heyden 1862 from Oligocene of Germany (28.4 to 23.03 Mya) (Von Heyden, 1862) are roughly concordant with our estimation, which dated splitting of *Dorcadion* from its last common ancestor genus *Morimu*s ∼26 Mya. The speciation event within the genus *Dorcadion* was 14.34 Mya in our estimation, earlier than the chronogram of Dascălu et al. (2022), who dated the beginning of speciation within the genera *Dorcadion* was around 9.8 Mya.

The emergence time of *Mesosa* was ∼67 Mya according to our chronogram, but ∼107 Mya according to Nie et al. (2020). All fossil records of *Mesosa* Latreille 1829 (syn. *Sinocalosoma* Hong 1983) were from the Miocene of China and the Oligocene of Germany (28.4 to 11.608 Mya), (Hong 1983, Hong and Wang 1986).

Considering our chronogram, the MRCA of four species of *Agapanthia* appeared about 55 Mya in Paleogene (Figure 2), while the MRCA of two *Agapanthia daurica* individuals dated 51.79 Mya in Nie at el. (2021). However, we could not access any fossil record for this genus.

The MRCA of *Pogonocherus* and *Exocentrus* emerged around 65 Mya at the edge of the Cretaceous and Paleogene, according to the chronogram of Ashman et al. (2022), while it is 37 Mya around mid-Paleogene, according to the hypothesized chronogram in the present study (Figure 2). The oldest fossil of *Pogonocherus, Pogonocherus jaekeli* Zang 1905, is from Baltic Amber, 37.2 to 33.9 Mya (Vitali, 2009).

We estimated the MRCA of Saperdini and Phytoeciini as 65 million years old, which the latter might be a synonym of the preceding in addition to Obereini (de Santana Souza et al. 2020). Ashman et al. (2022) estimated the MRCA of *Phytoecia* and *Oberea* 36 Mya. The oldest fossil of the genus *Saperda* is *Saperda caroli* Vitali 2015, from Uintan of Colorado, America (46.2 - 40.4 Mya) and the oldest fossil from Eurasia is *Saperda densipunctata* Théobald 1937, from Oligocene of Germany, 33.9 to 28.4 Mya (Vitali, 2015). The oldest Obereini fossil is the youngest in the group, *Oberea praemortua* von Heyden 1862, and from the Miocene of Germany, 15.97 to 11.608 Mya (Von Heyden, 1862).

According to our time estimation of the intrageneric diversification of the taxa involved in the analysis occurred during Cenozoic, mostly in Neogene, concurred with Ashman et al. (2022). Diversification of the members of *Agapanthia* seems to be occurred during Paleogene ∼55 Mya, earlier than *Phytoecia* and *Dorcadion* members, ∼23 Mya and ∼14 Mya, respectively, during Neogene.

What our study reveals is the Lamiinae specimens from East of Marmara Basin, Türkiye, were closely related to European conspecifics and congeners. And, relying on our samples, we stand by the side of de Santana Souza et al. (2020) in supporting the hypothesis of uniting the tribes Dorcadionini, Gnomini, Monochamini (in terms of *Monochamus*) and the current Lamiini under a single tribe Lamiini. Considering all recent studies and the present study, the emergence time of the subfamily Lamiinae is Jurassic to early Cretaceous, and the extant species included in this study presently occur in the East of Marmara Basin that emerged during the Neogene period.

## Supporting information

https://doi.org/10.6084/m9.figshare.21070867.v1

https://doi.org/10.6084/m9.figshare.21070825.v1

10.6084/m9.figshare.22864970

10.6084/m9.figshare.21070915

## Acknowledgements

We thank Dr Burcu Şabanoğlu Şimşek and Dr Hüseyin Özdikmen for their contributions to morphological identification, Dr Şener Atak and Osman Atak for their sampling efforts and Saadet Şirin Coşkun for visualisation of the figures. This study is a part of PhD thesis of Havva Kübra Soydabaş Ayoub and funded by the Scientific Research Foundation of Kocaeli University, grant number BAP 139-2018.

## Supplementary Materials

Supplementary Figure S1: The Neighbor-Joining tree of the subfamily Lamiinae comprises a representative haplotype for each available species from each zoogeographic region based on mitochondrial *COI* sequences (658 bp) obtained in this study and retrieved from the BOLD taxonomy archive

Supplementary Figure S2: The Maximum Likelihood tree of the subfamily Lamiinae comprises a representative haplotype for each available species from each zoogeographic region based on mitochondrial *COI* sequences (658 bp) obtained in this study and retrieved from the BOLD taxonomy archive

Supplementary Table S1. Binomial names, voucher codes, sampling coordinates, localities and GenBank accession numbers of the specimens sampled in this study Supplementary Table S2: Identity numbers, binomial names and sampling localities of mitochondrial *COI* gene region sequences of subfamily Lamiinae, which were retrieved from the BOLD taxonomy database.

## Data Availability Statement

All data generated or analyzed during this study are included in this published article [and its supplementary information files]. The sequences produced in this study are available in GenBank under the accession numbers OP279135-OP279183, OP279535-OP279581 and OP279486-OP279532.

