## Supplementary figures and images for "Time-scaled phylogenetic analysis of extant Lamiinae (Coleoptera, Cerambycidae) species of East of Marmara Basin, Türkiye, and their evolutionary similarity with the Eurasian congeners"

### https://doi.org/10.6084/m9.figshare.21070825.v1

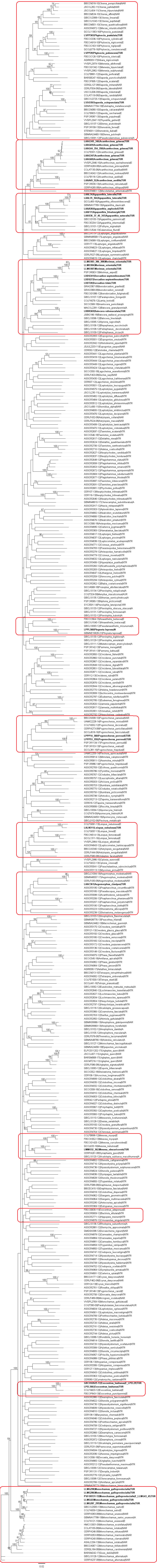

### https://doi.org/10.6084/m9.figshare.21070867.v1

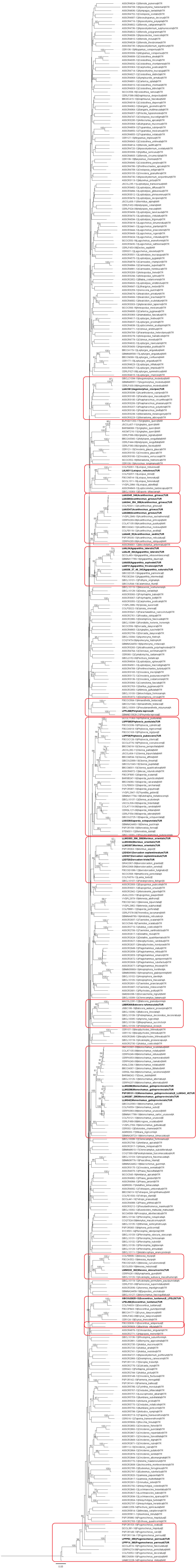
