## Supplementary material for "Time-scaled phylogenetic analysis of extant Lamiinae (Coleoptera, Cerambycidae) species of East of Marmara Basin, Türkiye, and their evolutionary similarity with the Eurasian congeners": 10.6084/m9.figshare.22864970

Table 1. Binomial names, voucher codes, sampling coordinates, localities and GenBank accession numbers of the specimens sampled in this study

| **#** | **Tribe** | **Species** | **Voucher ID** | **Coordinate** | **Locality** | **COI** | **16S rRNA** | **28S rRNA** |
| --- | --- | --- | --- | --- | --- | --- | --- | --- |
| 1 | Acanthocinini | *Acanthocinus* (Acanthocinus) *aedilis* (Linnaeus, 1758) | LAAA49 | 40°49'33.2"N 29°29'52.0"E | Timber Yard, Gebze, | OP279161 | - | - |
| 2 | Acanthocinini | *Acanthocinus* (Acanthocinus) *aedilis* (Linnaeus, 1758) | LAAA52 | 40°42'52.6"N 30°03'21.1"E | Wood Processing Plant, Kartepe | OP279162 | OP279556 | OP279504 |
| 3 | Acanthocinini | *Acanthocinus* (Acanthocinus) *griseus* (Fabricius, 1793) | LAAG345 | 40°49'01.6"N 29°29'49.6"E | Wood Processing Plant, Gebze | OP279165 | OP279562 | OP279509 |
| 4 | Acanthocinini | *Acanthocinus* (Acanthocinus) *griseus* (Fabricius, 1793) | LAAG346 | 40°49'33.3"N 29°29'52.1"E | Timber Yard, Gebze | OP279166 | OP279560 | OP279512 |
| 5 | Acanthocinini | *Acanthocinus* (Acanthocinus) *griseus* (Fabricius, 1793) | LAAG347 | 40°49'04.7"N 29°29'52.7"E | Timber Yard, Gebze | OP279163 | OP279561 | OP279507 |
| 6 | Acanthocinini | *Acanthocinus* (Acanthocinus) *griseus* (Fabricius, 1793) | LAAG354 | 40°49'45.4"N 29°55'05.4"E | Forest, İzmit | OP279168 | OP279559 | OP279508 |
| 7 | Acanthocinini | *Acanthocinus* (Acanthocinus) *griseus* (Fabricius, 1793) | LAAG358 | 40°41'31.4"N 29°53'55.2"E" | Countryside, Başiskele | OP279169 | - | OP279510 |
| 8 | Acanthocinini | *Acanthocinus* (Acanthocinus) *griseus* (Fabricius, 1793) | LAAG44 | 40°47'19.3"N 29°50'45.4"E | Forest, Derince | OP279167 | OP279557 | OP279506 |
| 9 | Acanthocinini | *Acanthocinus* (Acanthocinus) *griseus* (Fabricius, 1793) | LAAG68 | 40°49'39.8"N 29°29'47.6"E | Forest, Gebze | OP279164 | - | OP279505 |
| 10 | Acanthocinini | *Acanthocinus* (Acanthocinus) *griseus* (Fabricius, 1793) | LAAG69 | 40°49'05.6"N 29°31'08.0"E | Forest, Gebze | OP279170 | OP279558 | OP279511 |
| 11 | Acanthocinini | *Leiopus* (Leiopus) *nebulosus* (Linnaeus, 1758) | LALN311 | 40°49'28.1"N 29°29'48.5"E | Timber Yard, Gebze | OP279160 | OP279555 | OP279519 |
| 12 | Acanthoderini | *Aegomorphus clavipes* (Schrank, 1781) | LAAC33 | 40°49'30.5"N 29°30'02.3"E | Gebze Forest | - | OP279536 | OP279502 |
| 13 | Acanthoderini | *Aegomorphus clavipes* (Schrank, 1781) | LAAC361 | 40°49'45.1"N 29°29'50.9"E | Gebze Forest | OP279155 | OP279537 | OP279503 |
| 14 | Agapanthiini | *Agapanthia* (*Epoptes*) *asphodeli* (Latreille, 1804) | LAAA30 | 40°41'44.1"N 29°53'50.9"E | Başiskele, Countryside | OP279144 | OP279574 | OP279493 |
| 15 | Agapanthiini | *Agapanthia* (*Epoptes*) *lateralis* Ganglbauer, 1883 | LAAL34 | 40°41'49.4"N 29°53'40.9"E | Countryside, Başiskele | OP279141 | OP279575 | OP279494 |
| 16 | Agapanthiini | *Agapanthia* (*Epoptes*) *lateralis* Ganglbauer, 1883 | LAAL35 | 40°41'53.5"N 29°53'42.0"E | Başiskele, Countryside | OP279142 | OP279577 | OP279496 |
| 17 | Agapanthiini | *Agapanthia* (*Epoptes*) *lateralis* Ganglbauer, 1883 | LAAL364 | 40°41'49.2"N 29°54'06.2"E | Başiskele, Countryside | OP279143 | OP279576 | OP279495 |
| 18 | Agapanthiini | *Agapanthia* (Agapanthia) *suturalis* (Fabricius, 1787) | LAAS343 | 40°41'24.3"N 29°53'09.7"E | Başiskele Forest | OP279139 | OP279571 | OP279491 |
| 19 | Agapanthiini | *Agapanthia* (Agapanthia) *suturalis* (Fabricius, 1787) | LAAS36 | 40°42'01.4"N 29°54'24.8"E | Başiskele, Countryside | OP279136 | OP279573 | OP279492 |
| 20 | Agapanthiini | *Agapanthia* (Agapanthia) *suturalis* (Fabricius, 1787) | LAAS37 | 40°40'53.9"N 29°53'08.9"E | Başiskele, Countryside | OP279137 | OP279570 |  |
| 21 | Agapanthiini | *Agapanthia* (Agapanthia) *suturalis* (Fabricius, 1787) | LAAS46 | 40°41'44.4"N 29°48'54.8"E" | Gölcük, Countryside | OP279138 | OP279572 | OP279490 |
| 22 | Agapanthiini | *Agapanthia (Smaragdula) frivaldszkyi* Ganglbauer, 1884 | LAAF31 | 40°40'24.8"N 29°50'37.1"E | Gölcük, Countryside | OP279140 | OP279569 | OP279489 |
| 23 | Lamiini | *Morimus orientalis* Reitter, 1894 | LLMO303 | 41°07'53.0"N 30°11'55.6"E | Kandıra Forest | OP279173 | OP279546 | OP279529 |
| 24 | Lamiini | *Morimus orientalis* Reitter, 1894 | LLMO304 | 41°07'44.9"N 30°11'57.6"E | Kandıra Forest | OP279176 | OP279549 | OP279526 |
| 25 | Lamiini | *Morimus orientalis* Reitter, 1894 | LLMO306 | 40°49'45.1"N 29°29'50.9"E | Gebze Forest | OP279174 | OP279547 | OP279527 |
| 26 | Lamiini | *Morimus orientalis* Reitter, 1894 | LLMO307 | 40°41'20.7"N 29°54'05.7"E | Başiskele Countryside | OP279172 | OP279548 | OP279525 |
| 27 | Lamiini | *Morimus orientalis* Reitter, 1894 | LLMO308 | 40°49'07.8"N 29°53'54.1"E | Forest, İzmit | OP279175 | OP279551 |  |
| 28 | Lamiini | *Morimus orientalis* Reitter, 1894 | LLMO310 | 41°08'25.5"N 30°10'20.0"E | Forest, Kandıra | - | OP279550 | OP279528 |
| 29 | Monochamini | *Monochamus* (Monochamus) *galloprovincialis* (Olivier, 1795) | LMMG286 | 41°07'45.9"N 30°12'50.1"E | Forest, Kandıra | OP279148 | OP279540 | OP279523 |
| 30 | Monochamini | *Monochamus* (Monochamus) *galloprovincialis* (Olivier, 1795) | LMMG287 | 40°49'28.1"N 29°29'48.5"E | Timber Yard, Gebze | OP279153 | - | OP279520 |
| 31 | Monochamini | *Monochamus* (Monochamus) *galloprovincialis* (Olivier, 1795) | LMMG290 | 40°49'33.2"N 29°29'52.0"E | Timber Yard, Gebze | - | OP279542 | - |
| 32 | Monochamini | *Monochamus* (Monochamus) *galloprovincialis* (Olivier, 1795) | LMMG295 | 40°50'19.0"N 29°27'35.9"E | Timber Yard, Gebze | OP279154 | - | - |
| 33 | Monochamini | *Monochamus* (Monochamus) *galloprovincialis* (Olivier, 1795) | LMMG298 | 40°49'28.1"N 29°29'48.5"E | Timber Yard, Gebze | OP279151 | OP279543 | - |
| 34 | Monochamini | *Monochamus* (Monochamus) *galloprovincialis* (Olivier, 1795) | LMMG42 | 40°50'19.0"N 29°27'35.9"E | Timber Yard, Gebze | OP279152 | - | OP279521 |
| 35 | Monochamini | *Monochamus* (Monochamus) *galloprovincialis* (Olivier, 1795) | LMMG43 | 40°49'58.1"N 29°28'47.4"E | Timber Yard, Gebze | OP279149 | OP279539 | - |
| 36 | Monochamini | *Monochamus* (Monochamus) *galloprovincialis* (Olivier, 1795) | LMMG45 | 40°47'19.3"N 29°50'45.4"E | Forest, Derince | OP279150 | OP279541 | OP279522 |
| 37 | Dorcadionini | *Dorcadion (Cribridorcadion) septemlineatum* Waltl, 1838 | LDDS341 | 41°07'44.9"N 30°11'57.6"E | Forest, Kandıra | OP279182 | OP279553 | OP279531 |
| 38 | Dorcadionini | *Dorcadion (Cribridorcadion) septemlineatum* Waltl, 1838 | LDDS27 | 40°44'49.7"N 30°03'58.8"E | Forest, Kartepe | OP279183 | OP279554 | OP279530 |
| 39 | Dorcadionini | *Dorcadion (Maculatodorcadion) triste* Frivaldszky, 1845 | LDDT20 | 40°44'49.7"N 30°03'58.8"E | Forest, Kartepe | OP279181 | OP279552 | OP279532 |
| 40 | Mesosini | *Mesosa* (*Aplocnemia*) *obscuricornis* Pic, 1894 | LMMO32 | 40°41'57.5"N 29°54'28.6"E | Başiskele Countryside | OP279180 | OP279544 | OP279514 |
| 41 | Mesosini | *Mesosa* (*Aplocnemia*) *obscuricornis* Pic, 1894 | LMMO362 | 40°41'04.0"N 29°53'42.0"E | Başiskele Countryside | OP279179 | OP279545 | OP279513 |
| 42 | Pogonocherini | *Pogonocherus* (Pogonocherus) *perroudi* Mulsant, 1839 | LPPP14 | 40°49'45.1"N 29°29'50.9"E | Forest, Gebze | OP279156 | OP279568 | OP279517 |
| 43 | Pogonocherini | *Pogonocherus* (Pogonocherus) *perroudi* Mulsant, 1839 | LPPP53 | 40°40'23.1"N 30°04'08.1"E | Forest, Kartepe | OP279158 | OP279566 | OP279516 |
| 44 | Pogonocherini | *Pogonocherus* (Pogonocherus) *perroudi* Mulsant, 1839 | LPPP359 | 40°49'15.6"N 30°02'22.9"E | Forest, Kartepe | OP279159 | OP279567 | OP279515 |
| 45 | Pogonocherini | *Pogonocherus* (Pogonocherus) *perroudi* Mulsant, 1839 | LPPP360 | 40°57'19.0"N 29°38'54.4"E | Forest, Gebze | OP279157 | OP279565 | OP279518 |
| 46 | Pogonocherini | *Exocentrus* (Exocentrus) *lusitanus* (Linnaeus, 1767) | LAEL366 | 40°42'52.6"N 30°03'21.1"E | Timber Yard, Gebze | OP279145 | OP279563 | OP279486 |
| 47 | Pogonocherini | *Exocentrus* (Exocentrus) *lusitanus* (Linnaeus, 1767) | LAEL29 | 41°00'26.6"N 29°55'15.4"E | Forest, Gebze | OP279146 | OP279564 | OP279487 |
| 48 | Saperdini | *Saperda* (*Lopezcolonia*) *octopunctata* (Scopoli, 1772) | LSSO344 | 40°59'31.4"N 29°33'23.4"E | Forest, Gebze | - | - | OP279497 |
| 49 | Saperdini | *Saperda* (*Lopezcolonia*) *octopunctata* (Scopoli, 1772) | LSSO39 | 40°49'04.7"N 29°29'54.2"E | Timber Yard, Gebze | OP279171 | OP279578 | OP279498 |
| 50 | Saperdini | *Phytoecia (Helladia) praetextata* (Steven, 1817) | LPPP367 | 41°07'44.9"N 30°11'57.6"E | Kandıra Forest | - | OP279581 | OP279500 |
| 51 | Saperdini | *Phytoecia* (Phytoecia) *pustulata* (Schrank, 1776) | LPPP365 | 41°07'45.9"N 30°12'50.1"E | Kandıra Forest | OP279177 | OP279580 | OP279499 |
| 52 | Saperdini | *Phytoecia* (Phytoecia) *pubescens* Pic, 1895 | LPPP28 | 40°41'20.7"N 29°54'05.7"E | Başiskele Countryside | OP279178 | OP279579 | OP279501 |
| 53 | Batocerini | *Batocera rufomaculata* (DeGeer, 1775) | LBBR369 | n/a | Diyarbakır | OP279147 | OP279538 | OP279524 |
| 54 | Phrynetini | *Phryneta leprosa (Fabricius, 1775)* | LPPL368 | n/a | Cameroon (Intercepted in Port, Derince) | OP279135 | OP279535 | OP279488 |
