## Supplementary material for "Time-scaled phylogenetic analysis of extant Lamiinae (Coleoptera, Cerambycidae) species of East of Marmara Basin, Türkiye, and their evolutionary similarity with the Eurasian congeners": 10.6084/m9.figshare.21070915

Supplementary Table S1. Identity numbers, binomial names and sampling localities of mitochondrial COI gene region sequences of subfamily Lamiinae, which were retrieved from the BOLD taxonomy database

| **#** | **BOLD ID** | **Species** | **Locality** | **References** |
| --- | --- | --- | --- | --- |
| 1 | ASSCR2648-12 | *Plagiohammus elatus* | Costa Rica | Anonymous |
| 2 | ASSCR2901-12 | *Venustus zeteki* | Costa Rica | Anonymous |
| 3 | ASSCR2993-12 | *Eupogonius pubicollis* | Costa Rica | Anonymous |
| 4 | ASSCR3303-12 | *Apteralcidion lapierrei* | Costa Rica | Anonymous |
| 5 | ASSCR4575-12 | *Leptostylus pygialis* | Costa Rica | Anonymous |
| 6 | ASSCR4637-12 | *Lepturges proxima* | Costa Rica | Anonymous |
| 7 | ASSCR4758-12 | *Oedopeza setigera* | Costa Rica | Anonymous |
| 8 | ASSCR4822-12 | *Stenolis polygramma* | Costa Rica | Anonymous |
| 9 | CERLF578-08 | *Threnetica lacrymans* | Thailand | Anonymous |
| 10 | CERPA261-08 | *Synaphaeta guexi* | Canada | Anonymous |
| 11 | CERPA383-08 | *Monochamus urussovi* | Russia | Anonymous |
| 12 | FBCOC163-10 | *Phytoecia nigripes* | Germany | Anonymous |
| 13 | GBCOU2725-13 | *Saperda octopunctata* | Germany | Hendrich et al. (2015) |
| 14 | GBCL10132-12 | *Pterolophia kubokii* | Taiwan | Yeh et al. (Unpublished) |
| 15 | GBMNA28723-19 | *Monochamus bimaculatus* | India | Behere ve diğ., (Unpublished) |
| 16 | SICOC545-18 | *Amillarus apicalis* | Nicaragua | Anonymous |
| 17 | BARSM068-17 | *Urgleptes querci* | Canada | Dewaard (Unpublished) |
| 18 | ASSCR2710-12 | *Adetus costicollis* | Costa Rica | Anonymous |
| 19 | ASSCR2917-12 | *Ataxia nivisparsa* | Costa Rica | Anonymous |
| 20 | ASSCR3444-12 | *Lagocheirus praecellens* | Costa Rica | Anonymous |
| 21 | ASSCR4604-12 | *Leptostylus xgriseus* | Costa Rica | Anonymous |
| 22 | ASSCR4876-12 | *Omosarotes singularis* | Costa Rica | Anonymous |
| 23 | GBCL10105-12 | *Anoplophora macularia* | Taiwan | Yeh et al. (Unpublished) |
| 24 | GBMNA17794-19 | *Eutetrapha metallescens* | China | Yang (Unpublished) |
| 25 | GBMNA24459-19 | *Apomecyna cretacea* | India | Behere et al. (Unpublished) |
| 26 | GBMNA24464-19 | *Batocera lineolata* | India | Behere et al. (Unpublished) |
| 27 | ASSCR3424-12 | *Lagocheirus cristulatus* | Costa Rica | Anonymous |
| 28 | CER112-13 | *Oncideres rubra* | Mexico | Anonymous |
| 29 | ASSCR2672-12 | *Plagiohammus spinipennis* | Costa Rica | Anonymous |
| 30 | ASSCR2851-12 | *Oncideres fulvostillata* | Costa Rica | Anonymous |
| 31 | ASSCR2860-12 | *Oncideres punctata* | Costa Rica | Anonymous |
| 32 | ASSCR2936-12 | *Aerenea impetiginosa* | Costa Rica | Anonymous |
| 33 | ASSCR2980-12 | *Desmiphora fasciculata* | Costa Rica | Anonymous |
| 34 | ASSCR3259-12 | *Xenocona pulchra* | Costa Rica | Anonymous |
| 35 | ASSCR3399-12 | *Hylettus seniculus* | Costa Rica | Anonymous |
| 36 | ASSCR4771-12 | *Ozineus arietinus* | Costa Rica | Anonymous |
| 37 | ASSCR5750-12 | *Itumbiara picticornis* | Costa Rica | Anonymous |
| 38 | FBCOG1425-12 | *Mesosa curculionoides* | France | Anonymous |
| 39 | JSCOL282-11 | *Oberea pallida* | Canada | Hebert et al. (2016) |
| 40 | CER121-13 | *Oreodera glauca glauca* | Mexico | Anonymous |
| 41 | GBCL10130-12 | *Pterolophia bigibbera* | Taiwan | Yeh et al. (Unpublished) |
| 42 | VVGPL2984-15 | *Leiopus albivittis* | Russia | Grebennikov et al. (2017) |
| 43 | ASSCR2766-12 | *Amillarus apicalis* | Costa Rica | Anonymous |
| 44 | ASSCR3223-12 | *Steirastoma albiceps* | Costa Rica | Anonymous |
| 45 | ASSCR3499-12 | *Leptostylus laevicauda* | Costa Rica | Anonymous |
| 46 | CER007-12 | *Lagocheirus obsoletus* | Mexico | Anonymous |
| 47 | FBCOD630-11 | *Exocentrus adspersus* | Germany | Hendrich et al. (2015) |
| 48 | ASSCR2808-12 | *Ischiocentra monteverdensis* | Costa Rica | Anonymous |
| 49 | ASSCR3406-12 | *Hyperplatys pusillus* | Costa Rica | Anonymous |
| 50 | CERPA282-08 | *Anoplophora glabripennis* | USA | Anonymous |
| 51 | FBCOK552-13 | *Mesosa myops* | Mongolia | Hendrich et al. (2015) |
| 52 | GBCOU2583-13 | *Monochamus sartor* | Austria | Hendrich et al. (2015) |
| 53 | GCOL483-16 | *Agapanthia villosoviridescens* | Germany | Rulik et al. (2017) |
| 54 | ASSCR4747-12 | *Oedopeza leucostigma* | Costa Rica | Anonymous |
| 55 | ASSCR4982-12 | *Phaea giesberti* | Costa Rica | Anonymous |
| 56 | CERGL127-08 | *Saperda imitans* | Canada | Anonymous |
| 57 | JSCOL404-11 | *Oberea tripunctata* | Canada | Hebert et al. (2016) |
| 58 | PSFOR148-13 | *Pogonocherus caroli* | France | Rougerie et al. (2015) |
| 59 | GBCL10101-12 | *Pothyne virginalis* | Taiwan | Yeh et al. (Unpublished) |
| 60 | GBCL10107-12 | *Paraleprodera itzingeri* | Taiwan | Yeh et al. (Unpublished) |
| 61 | ASCMT216-11 | *Urgleptes querci* | Canada | Dewaard (Unpublished) |
| 62 | ASSCR2624-12 | *Deliathis quadritaeniator* | Costa Rica | Anonymous |
| 63 | ASSCR2637-12 | *Neoptychodes hondurae* | Costa Rica | Anonymous |
| 64 | ASSCR2677-12 | *Plagiohammus thiodes* | Costa Rica | Anonymous |
| 65 | ASSCR2691-12 | *Taeniotes praeclarus* | Costa Rica | Anonymous |
| 66 | ASSCR2741-12 | *Adetus nesiotes* | Costa Rica | Anonymous |
| 67 | ASSCR2785-12 | *Eudesmus ferrugineus* | Costa Rica | Anonymous |
| 68 | GBMNA17788-19 | *Monochamus sartor urussovii* | China | Yang (Unpublished) |
| 69 | ASSCR2820-12 | *Jamesia papulenta* | Costa Rica | Anonymous |
| 70 | ASSCR3001-12 | *Eupogonius ursulus* | Costa Rica | Anonymous |
| 71 | ASSCR3148-12 | *Oreodera fluctuosa* | Costa Rica | Anonymous |
| 72 | ASSCR3180-12 | *Oreodera verrucosa* | Costa Rica | Anonymous |
| 73 | ASSCR3470-12 | *Leptostylus cristulatus* | Costa Rica | Anonymous |
| 74 | ASSCR3476-12 | *Leptostylus decipiens* | Costa Rica | Anonymous |
| 75 | ASSCR4615-12 | *Lepturges charicles* | Costa Rica | Anonymous |
| 76 | ASSCR4647-12 | *Lithargyrus melzeri* | Costa Rica | Anonymous |
| 77 | ASSCR4699-12 | *Nyssodectes roseicollis* | Costa Rica | Anonymous |
| 78 | ASSCR4779-12 | *Ozineus moestus* | Costa Rica | Anonymous |
| 79 | ASSCR4810-12 | *Stenolis inclusa* | Costa Rica | Anonymous |
| 80 | ASSCR4828-12 | *Stenolis theobromae* | Costa Rica | Anonymous |
| 81 | ASSCR4838-12 | *Leptocometes acutispinis* | Costa Rica | Anonymous |
| 82 | ASSCR4881-12 | *Carneades hemileuca* | Costa Rica | Anonymous |
| 83 | ASSCR4943-12 | *Colobothea unilineata* | Costa Rica | Anonymous |
| 84 | ASSCR4957-12 | *Priscilla hypsiomoides* | Costa Rica | Anonymous |
| 85 | ASSCR4972-12 | *Mecas rotundicollis* | Costa Rica | Anonymous |
| 86 | ASSCR4988-12 | *Phaea phthisica* | Costa Rica | Anonymous |
| 87 | ASSCR5757-12 | *Leucophoebe albaria* | Costa Rica | Anonymous |
| 88 | ASSCR5806-12 | *Recchia hirsuta* | Costa Rica | Anonymous |
| 89 | CER080-12 | *Cymatonycha castanea* | Mexico | Anonymous |
| 90 | COLFB118-12 | *Acanthocinus aedilis* | Finland | Pentinsaari et al. (2014) |
| 91 | COLFB898-12 | *Mesosa myops* | Finland | Pentinsaari et al. (2014) |
| 92 | COQT504-09 | *Parahybolasias fuscomaculatus* | Australia | Anonymous |
| 93 | FBCOE204-12 | *Agapanthia intermedia* | Germany | Hendrich et al. (2015) |
| 94 | FBCOF895-12 | *Saperda scalaris* | Germany | Hendrich et al. (2015) |
| 95 | FBCOG1086-12 | *Iberodorcadion fuliginator* | Germany | Anonymous |
| 96 | GBCOU2625-13 | *Exocentrus lusitanus* | Slovenia | Hendrich et al. (2015) |
| 97 | JSCOL248-11 | *Oberea perspicillata* | Canada | Hebert et al. (2016) |
| 98 | JSCOL406-11 | *Sternidius alpha* | Canada | Hebert et al. (2016) |
| 99 | PSFOR063-13 | *Morimus asper* | France | Rougerie et al. (2015) |
| 100 | PSFOR065-13 | *Niphona picticornis* | France | Rougerie et al. (2015) |
| 101 | CERLF430-08 | *Aegomorphus modestus* | Canada | Anonymous |
| 102 | CERLF462-08 | *Ecyrus dasycerus* | Canada | Anonymous |
| 103 | CERLF531-08 | *Psenocerus supernotatus* | Canada | Anonymous |
| 104 | VVGPL2769-15 | *Monochamus guttulatus* | Russia | Grebennikov et al. (2017) |
| 105 | VVGPL2947-15 | *Thyestilla gebleri* | Russia | Grebennikov et al. (2017) |
| 106 | VVGPL2982-15 | *Menesia sulphurata* | Russia | Grebennikov et al. (2017) |
| 107 | CERPA240-08 | *Monochamus clamator* | Canada | Anonymous |
| 108 | FBCOP843-13 | *Exocentrus punctipennis* | Slovenia | Hendrich et al. (2015) |
| 109 | GBCOU2999-13 | *Oberea linearis* | Germany | Hendrich et al. (2015) |
| 110 | GBCL10083-12 | *Eoporis bifasciana* | Taiwan | Yeh et al. (Unpublished) |
| 111 | GBCL10094-12 | *Apomecyna histrio* | Taiwan | Yeh et al. (Unpublished) |
| 112 | GBCL10102-12 | *Anoplophora davidis* | Taiwan | Yeh et al. (Unpublished) |
| 113 | GBCL10103-12 | *Anoplophora flavomaculata* | Taiwan | Yeh et al. (Unpublished) |
| 114 | GBCL10106-12 | *Blepephaeus succinctor* | Taiwan | Yeh et al. (Unpublished) |
| 115 | GBCL10109-12 | *Peblephaeus ziczac* | Taiwan | Yeh et al. (Unpublished) |
| 116 | GBCL10118-12 | *Acalolepta grossescapus* | Taiwan | Yeh et al. (Unpublished) |
| 117 | GBCL10126-12 | *Monochamus alternatus* | Taiwan | Yeh et al. (Unpublished) |
| 118 | GBCL10127-12 | *Monochamus fascioguttatus* | Taiwan | Yeh et al. (Unpublished) |
| 119 | GBCL10128-12 | *Desisa variabilis* | Taiwan | Yeh et al. (Unpublished) |
| 120 | GBCL10131-12 | *Pterolophia formosana* | Taiwan | Yeh et al. (Unpublished) |
| 121 | GBCL10137-12 | *Glenea acutoides* | Taiwan | Yeh et al. (Unpublished) |
| 122 | GBCL10138-12 | *Xenolea asiatica* | Taiwan | Yeh et al. (Unpublished) |
| 123 | GBMIN48610-17 | *Olenecamptus subobliteratus* | China | Anonymous |
| 124 | GBMNA17789-19 | *Agapanthia daurica* | China | Anonymous |
| 125 | SICOC546-18 | *Taeniotes scalatus* | Nicaragua | Anonymous |
| 126 | SICOC558-18 | *Colobothea ramosa* | Nicaragua | Anonymous |
| 127 | BARSM242-17 | *Goes debilis* | Canada | Dewaard (Unpublished) |
| 128 | GBCLC1094-19 | *Aegomorphus modestus* | USA | Caterino ve Kadau (Unpublished) |
| 129 | JSBIC185-18 | *Batocera laena* | Papua New Guinea | Anonymous |
| 130 | GBMNA24462-19 | *Macrochenus guerinii* | India | Behere et al. (Unpublished) |
| 131 | AGIRI090-17 | *Astathes bimaculata* | India | Anonymous |
| 132 | GBCL0152-06 | *Plectrura metallica* | Japan | Nakamine (Unpublished) |
| 133 | ASSCR2763-12 | *Sarillus pygmaeus* | Costa Rica | Anonymous |
| 134 | ASSCR2827-12 | *Lochmaeocles batesi* | Costa Rica | Anonymous |
| 135 | ASSCR3175-12 | *Oreodera semialba* | Costa Rica | Anonymous |
| 136 | ASSCR3207-12 | *Psapharochrus polysticta* | Costa Rica | Anonymous |
| 137 | ASSCR3294-12 | *Anisopodus xylinus* | Costa Rica | Anonymous |
| 138 | ASSCR3306-12 | *Atrypanius conspersus* | Costa Rica | Anonymous |
| 139 | ASSCR3363-12 | *Carphontes posticalis* | Costa Rica | Anonymous |
| 140 | ASSCR3366-12 | *Cosmotoma fasciata* | Costa Rica | Anonymous |
| 141 | ASSCR4623-12 | *Lepturges infilatus* | Costa Rica | Anonymous |
| 142 | ASSCR4632-12 | *Lepturges navicularisAS3* | Costa Rica | Anonymous |
| 143 | ASSCR4735-12 | *Nyssodrysternum signiferum* | Costa Rica | Anonymous |
| 144 | ASSCR4836-12 | *Sympagus laetabilis* | Costa Rica | Anonymous |
| 145 | ASSCR4932-12 | *Colobothea chontalensis* | Costa Rica | Anonymous |
| 146 | ASSCR4964-12 | *Sangaris multimaculata* | Costa Rica | Anonymous |
| 147 | BBCCN013-10 | *Tetraopes tetrophthalmus* | Canada | Hebert et al. (2016) |
| 148 | BBCEC415-10 | *Graphisurus fasciatus* | Canada | Hebert et al. (2016) |
| 149 | FBCOG1342-12 | *Menesia bipunctata* | Germany | Hendrich et al. (2015) |
| 150 | GBCOD105-13 | *Agapanthia pannonica* | Germany | Hendrich et al. (2015) |
| 151 | GBCOU1134-13 | *Lamia textor* | Germany | Hendrich et al. (2015) |
| 152 | PSFOR158-13 | *Stenostola ferrea* | France | Rougerie et al. (2015) |
| 153 | CERPA353-08 | *Monochamus marmorator* | Canada | Anonymous |
| 154 | GRACI368-08 | *Iberodorcadion zenete* | Spain | Anonymous |
| 155 | GBCL10095-12 | *Apriona rugicollis* | Taiwan | Yeh et al. (Unpublished) |
| 156 | GBCL10099-12 | *Olenecamptus Taiwanus* | Taiwan | Yeh et al. (Unpublished) |
| 157 | GBCL10100-12 | *Moechotypa formosana* | Taiwan | Yeh et al. (Unpublished) |
| 158 | GBCL10108-12 | *Peblephaeus decoloratus decoloratus* | Taiwan | Yeh et al. (Unpublished) |
| 159 | GBCL10110-12 | *Mesoereis koshunensis* | Taiwan | Yeh et al. (Unpublished) |
| 160 | GBCL10111-12 | *Mutatocoptops anancyloides* | Taiwan | Yeh et al. (Unpublished) |
| 161 | GBCL10133-12 | *Pterolophia obscura obscura* | Taiwan | Yeh et al. (Unpublished) |
| 162 | GBMNA48780-19 | *Aristobia reticulator* | India | Anonymous |
| 163 | GBCLC1104-19 | *Batocera lineolata* | China | Anonymous |
| 164 | GBMNA24458-19 | *Epepeotes uncinatus* | India | Anonymous |
| 165 | GBMNA24460-19 | *Glenea pulchra* | India | Anonymous |
| 166 | SICOC566-18 | *Anisopodus mexicanus* | Nicaragua | Anonymous |
| 167 | STEN001-12 | *Stenostola dubia* | Norway | Kvamme et al. (2012) |
| 168 | GCOL633-16 | *Mesosa nebulosa* | Germany | Rulik et al. (2017) |
| 169 | ASSCR2669-12 | *Plagiohammus rubefactus* | Costa Rica | Anonymous |
| 170 | ASSCR2687-12 | *Taeniotes iridescens* | Costa Rica | Anonymous |
| 171 | ASSCR2756-12 | *Dorcasta dasycera* | Costa Rica | Anonymous |
| 172 | ASSCR2776-12 | *Cylicasta nysaAS3* | Costa Rica | Anonymous |
| 173 | ASSCR2871-12 | *Oncideres rubra* | Costa Rica | Anonymous |
| 174 | ASSCR3155-12 | *Oreodera granulifera* | Costa Rica | Anonymous |
| 175 | ASSCR3206-12 | *Psapharochrus phasianus* | Costa Rica | Anonymous |
| 176 | GCOL11363-16 | *Phytoecia pustulata* | Germany | Rulik et al. (2017) |
| 177 | SICOC550-18 | *Lagocheirus araneiformis* | Nicaragua | Anonymous |
| 178 | SICOC552-18 | *Steirastoma histrionica* | Nicaragua | Anonymous |
| 179 | ASSCR3190-12 | *Psapharochrus bivitta* | Costa Rica | Anonymous |
| 180 | ASSCR3369-12 | *Eutrypanus mucoreus* | Costa Rica | Anonymous |
| 181 | ASSCR3383-12 | *Hamatastus fasciatus* | Costa Rica | Anonymous |
| 182 | ASSCR3501-12 | *Leptostylus leucopygus* | Costa Rica | Anonymous |
| 183 | ASSCR4600-12 | *Leptostylus viriditinctus* | Costa Rica | Anonymous |
| 184 | ASSCR4682-12 | *Nealcidion scutellatum* | Costa Rica | Anonymous |
| 185 | ASSCR4708-12 | *Nyssodrysina haldemani* | Costa Rica | Anonymous |
| 186 | ASSCR4786-12 | *Pentheochaetes apicalis* | Costa Rica | Anonymous |
| 187 | ASSCR4843-12 | *Leptocometes barbiscapus* | Costa Rica | Anonymous |
| 188 | ASSCR4855-12 | *Trypanidius notatus* | Costa Rica | Anonymous |
| 189 | ASSCR4878-12 | *Carneades championi* | Costa Rica | Anonymous |
| 190 | ASSCR4969-12 | *Sympleurotis armatus* | Costa Rica | Anonymous |
| 191 | ASSCR4992-12 | *Tetraopes umbonatus* | Costa Rica | Anonymous |
| 192 | ASSCR5003-12 | *Arixiuna prolixa* | Costa Rica | Anonymous |
| 193 | ASSCR5753-12 | *Itumbiara subdilatata* | Costa Rica | Anonymous |
| 194 | ASSCR5767-12 | *Oedudes bifasciata* | Costa Rica | Anonymous |
| 195 | ASSCR5798-12 | *Antodice nympha* | Costa Rica | Anonymous |
| 196 | BBCCA042-12 | *Acanthocinus nodosus* | USA | Anonymous |
| 197 | BBCCM534-10 | *Oberea affinis* | Canada | Anonymous |
| 198 | CER092-12 | *Estoloides chamelae* | Mexico | Anonymous |
| 199 | CERLF457-08 | *Dorcaschema nigrum* | Canada | Anonymous |
| 200 | CERLF624-08 | *Astylopsis macula* | Canada | Anonymous |
| 201 | GBCOU1540-13 | *Anaesthetis testacea* | Italy | Hendrich et al. (2015) |
| 202 | GMGSK098-12 | *Dorcaschema nigrum* | USA | Anonymous |
| 203 | UAMIC2228-14 | *Pogonocherus mixtus* | USA | Sikes et al. (2016) |
| 204 | VVGPL2986-15 | *Oplosia suvorovi* | Russia | Grebennikov et al. (2017) |
| 205 | SYC3591-14 | *Pterolophia lateripicta* | French Polynesia | Ramage et al. (Unpublished) |
| 206 | GBCL10120-12 | *Acalolepta sublusca maculihumera* | Taiwan | Yeh et al. (Unpublished) |
| 207 | GBCL10136-12 | *Rhodopina subuniformis* | Taiwan | Yeh et al. (Unpublished) |
| 208 | GCOL381-16 | *Pogonocherus hispidus* | Germany | Rulik et al. (2017) |
| 209 | GCOL589-16 | *Iberodorcadion fuliginator* | Germany | Rulik et al. (2017) |
| 210 | GBMIN48557-17 | *Aegomorphus modestus* | USA | Caterino ve Kadau (Unpublished) |
| 211 | GCOL441-16 | *Tetrops praeustus* | Germany | Rulik et al. (2017) |
| 212 | AGIRI093-17 | *Stibara nigricornis* | India | Anonymous |
| 213 | CER136-13 | *Atrypanius conspersus* | Mexico | Anonymous |
| 214 | GCOL4516-16 | *Pogonocherus fasciculatus* | Germany | Rulik et al. (2017) |
| 215 | SICOA568-18 | *Prosoplus albofasciatus* | Papua New Guinea | Anonymous |
| 216 | ASSCR2658-12 | *Plagiohammus nitidus* | Costa Rica | Anonymous |
| 217 | ASSCR2697-12 | *Taeniotes scalarisAS3* | Costa Rica | Anonymous |
| 218 | ASSCR2735-12 | *Adetus mucoreus* | Costa Rica | Anonymous |
| 219 | ASSCR2744-12 | *Adetus pictus* | Costa Rica | Anonymous |
| 220 | ASSCR2783-12 | *Ecthoea quadricornis* | Costa Rica | Anonymous |
| 221 | ASSCR2787-12 | *Eudesmus rubefactus* | Costa Rica | Anonymous |
| 222 | ASSCR2804-12 | *Hesychotypa turbida* | Costa Rica | Anonymous |
| 223 | ASSCR2844-12 | *Oncideres albomarginata* | Costa Rica | Anonymous |
| 224 | ASSCR2849-12 | *Oncideres digna* | Costa Rica | Anonymous |
| 225 | ASSCR2942-12 | *Atelodesmis piperita* | Costa Rica | Anonymous |
| 226 | ASSCR3041-12 | *Parachalastinus rubrocinctus* | Costa Rica | Anonymous |
| 227 | ASSCR3067-12 | *Polyrhaphis belti* | Costa Rica | Anonymous |
| 228 | ASSCR3098-12 | *Acanthoderes rubripes* | Costa Rica | Anonymous |
| 229 | ASSCR3165-12 | *Oreodera inscripta* | Costa Rica | Anonymous |
| 230 | ASSCR3172-12 | *Oreodera purpurascens* | Costa Rica | Anonymous |
| 231 | ASSCR3195-12 | *Psapharochrus circumflexa* | Costa Rica | Anonymous |
| 232 | ASSCR3418-12 | *Lagocheirus binumeratus* | Costa Rica | Anonymous |
| 233 | ASSCR3430-12 | *Lagocheirus kathleenae* | Costa Rica | Anonymous |
| 234 | ASSCR3482-12 | *Leptostylus diffusus* | Costa Rica | Anonymous |
| 235 | ASSCR3484-12 | *Leptostylus gibbulosus* | Costa Rica | Anonymous |
| 236 | ASSCR3512-12 | *Leptostylus phrissominus* | Costa Rica | Anonymous |
| 237 | ASSCR4561-12 | *Leptostylus macrostigma* | Costa Rica | Anonymous |
| 238 | ASSCR4627-12 | *Lepturges limpidus* | Costa Rica | Anonymous |
| 239 | ASSCR4661-12 | *Mecotetartus antennatus* | Costa Rica | Anonymous |
| 240 | ASSCR4676-12 | *Nealcidion privatum* | Costa Rica | Anonymous |
| 241 | ASSCR4714-12 | *Nyssodrysina leucopyga* | Costa Rica | Anonymous |
| 242 | ASSCR4727-12 | *Nyssodrysternum poriferum* | Costa Rica | Anonymous |
| 243 | ASSCR4736-12 | *Nyssodrysternum sulphurescens* | Costa Rica | Anonymous |
| 244 | ASSCR4783-12 | *Paranisopodus heterotarsus* | Costa Rica | Anonymous |
| 245 | ASSCR4802-12 | *Stenolis calligramma* | Costa Rica | Anonymous |
| 246 | ASSCR4805-12 | *Stenolis circumscripta* | Costa Rica | Anonymous |
| 247 | ASSCR4824-12 | *Stenolis pulverea* | Costa Rica | Anonymous |
| 248 | ASSCR4850-12 | *Trypanidius mexicanus* | Costa Rica | Anonymous |
| 249 | ASSCR4859-12 | *Trypanidius rubripes* | Costa Rica | Anonymous |
| 250 | ASSCR4865-12 | *Urgleptes kuscheli* | Costa Rica | Anonymous |
| 251 | ASSCR4884-12 | *Carneades superba* | Costa Rica | Anonymous |
| 252 | ASSCR4895-12 | *Carterica pygmaea* | Costa Rica | Anonymous |
| 253 | ASSCR4918-12 | *Colobothea dispersa* | Costa Rica | Anonymous |
| 254 | ASSCR4962-12 | *Sangaris geometrica* | Costa Rica | Anonymous |
| 255 | ASSCR5770-12 | *Oedudes notaticollis* | Costa Rica | Anonymous |
| 256 | BBCCA067-12 | *Monochamus titillator* | USA | Anonymous |
| 257 | BBCCA179-12 | *Lepturges angulatus* | USA | Anonymous |
| 258 | BBCCN068-10 | *Lepturges confluens* | Canada | Hebert et al. (2016) |
| 259 | CER042-12 | *Peritapnia pilosa* | Mexico | Anonymous |
| 260 | COLFA121-10 | *Monochamus urussovii* | Finland | Pentinsaari et al. (2014) |
| 261 | COLFA552-12 | *Pogonocherus decoratus* | Finland | Pentinsaari et al. (2014) |
| 262 | COLFB881-12 | *Saperda perforata* | Finland | Pentinsaari et al. (2014) |
| 263 | COQT324-09 | *Menyllus maculicornus* | Australia | Anonymous |
| 264 | COQT390-09 | *Parahybolasias fuscomaculatus* | Australia | Anonymous |
| 265 | FBCOC036-10 | *Phytoecia cylindrica* | Germany | Anonymous |
| 266 | PSFOR142-13 | *Parmena meregallii* | France | Rougerie, et al. (2015) |
| 267 | PSFOR153-13 | *Pogonocherus ovatus* | France | Rougerie, et al. (2015) |
| 268 | CERGL184-08 | *Monochamus carolinensis* | Canada | Anonymous |
| 269 | GBCOU544-13 | *Calamobius filum* | Greece | Hendrich et al. (2015) |
| 270 | UAMIC2259-14 | *Plectrura spinicauda* | USA | Sikes et al. (2016) |
| 271 | VVGPL2979-15 | *Menesia albifrons* | Russia | Grebennikov et al. (2017) |
| 272 | CERLF580-08 | *Urgleptes facetus* | Canada | Anonymous |
| 273 | CERLF627-08 | *Lepturges symmetricus* | Canada | Anonymous |
| 274 | CERPA237-08 | *Monochamus alternatus* | Canada | Anonymous |
| 275 | CERPA265-08 | *Acanthocinus obliquus* | Canada | Anonymous |
| 276 | CERPA269-08 | *Acanthocinus princeps* | Canada | Anonymous |
| 277 | CERPA273-08 | *Pogonocherus penicillatus* | Canada | Anonymous |
| 278 | COLAT105-08 | *Monochamus notatus* | Canada | Anonymous |
| 279 | COLAT135-08 | *Acanthocinus pusillus* | Canada | Anonymous |
| 280 | GRACI367-08 | *Iberodorcadion graellsii* | Spain | Anonymous |
| 281 | GBCL10098-12 | *Olenecamptus formosanus* | Taiwan | Yeh et al. (Unpublished) |
| 282 | GBCL10119-12 | *Acalolepta permutans paucipunctata* | Taiwan | Yeh et al. (Unpublished) |
| 283 | GBCL10129-12 | *Pterolophia annulata* | Taiwan | Yeh et al. (Unpublished) |
| 284 | GBCL10134-12 | *Pterolophia reduplicata* | Taiwan | Yeh et al. (Unpublished) |
| 285 | GCOL6776-16 | *Phytoecia coerulescens* | Czech Republic | Rulik et al. (2017) |
| 286 | SICOC568-18 | *Imantocera penicillata* | Nepal | Anonymous |
| 287 | CER107-13 | *Neoptychodes trilineatus* | Mexico | Anonymous |
| 288 | CER138-13 | *Baryssinus chemsaki* | Mexico | Anonymous |
| 289 | JSBIC186-18 | *Batocera wallacei proserpina* | Papua New Guinea | Anonymous |
| 290 | ELPCG11202-17 | *Urgleptes querci* | Canada | Anonymous |
| 291 | GCOL1311-16 | *Leiopus femoratus* | Germany | Rulik et al. (2017) |
| 292 | ASSCR2653-12 | *Plagiohammus emanon* | Costa Rica | Anonymous |
| 293 | ASSCR2681-12 | *Ptychodes politus* | Costa Rica | Anonymous |
| 294 | ASSCR2720-12 | *Adetus analis* | Costa Rica | Anonymous |
| 295 | ASSCR2771-12 | *Hippopsis meinerti* | Costa Rica | Anonymous |
| 296 | ASSCR2797-12 | *Hesychotypa heraldica* | Costa Rica | Anonymous |
| 297 | ASSCR3278-12 | *Anisopodus hamaticollis* | Costa Rica | Anonymous |
| 298 | ASSCR3283-12 | *Anisopodus hiekei* | Costa Rica | Anonymous |
| 299 | ASSCR3350-12 | *Carphontes posticalis* | Costa Rica | Anonymous |
| 300 | ASSCR4720-12 | *Nyssodrysternum ocellatum* | Costa Rica | Anonymous |
| 301 | ASSCR4752-12 | *Oedopeza ocellator* | Costa Rica | Anonymous |
| 302 | ASSCR4891-12 | *Carterica optata* | Costa Rica | Anonymous |
| 303 | ASSCR4984-12 | *Phaea janzeni* | Costa Rica | Anonymous |
| 304 | ASSCR4903-12 | *Colobothea bitincta* | Costa Rica | Anonymous |
| 305 | CERLF432-08 | *Astylopsis collaris* | Canada | Anonymous |
| 306 | CERLF489-08 | *Microgoes oculatus* | Canada | Anonymous |
| 307 | CERLF588-08 | *Graphisurus despectus* | Canada | Anonymous |
| 308 | COLFG057-13 | *Leiopus linnei* | Estonia | Pentinsaari et al. (2014) |
| 309 | COQT479-09 | *Apomecyna histrio* | Australia | Anonymous |
| 310 | FBCON514-13 | *Leiopus femoratus* | Germany | Hendrich et al. (2015) |
| 311 | GBCOB007-12 | *Exocentrus adspersus* | Germany | Hendrich et al. (2015) |
| 312 | PSFOR087-13 | *Saperda populnea* | France | Rougerie, et al. (2015) |
| 313 | PSFOR141-13 | *Parmena balteus* | France | Rougerie, et al. (2015) |
| 314 | USCOL506-09 | *Saperda tridentata* | USA | Anonymous |
| 315 | CER116-13 | *Neoptychodes trilineatus* | Mexico | Anonymous |
| 316 | GBCL10091-12 | *Pseudocalamobius pubescens* | Taiwan | Yeh et al. (Unpublished) |
| 317 | GBCL10093-12 | *Pseudanaesthetis mizunumai* | Taiwan | Yeh et al. (Unpublished) |
| 318 | GBCL10135-12 | *Sthenias semicylindricus* | Taiwan | Yeh et al. (Unpublished) |
| 319 | GBMIN38776-13 | *Psacothea hilaris* | Italy | Hendrich et al. (2015) |
| 320 | BARSB247-16 | *Saperda puncticollis* | Canada | Anonymous |
| 321 | PSFOR1131-17 | *Deroplia troberti* | Morocco | Rougerie, et al. (2015) |
| 322 | SICOC556-18 | *Dorcasta dasycera* | Nicaragua | Anonymous |
| 323 | ASALC294-13 | *Monochamus notatus* | Canada | Anonymous |
| 324 | ASSCR2617-12 | *Deliathis nivea* | Costa Rica | Anonymous |
| 325 | ASSCR2627-12 | *Neoptychodes candidus* | Costa Rica | Anonymous |
| 326 | ASSCR2706-12 | *Taeniotes xanthostictus* | Costa Rica | Anonymous |
| 327 | ASSCR2770-12 | *Helvina howdenorum* | Costa Rica | Anonymous |
| 328 | ASSCR2817-12 | *Jamesia multivittata* | Costa Rica | Anonymous |
| 329 | ASSCR2840-12 | *Lochmaeocles tessellatus* | Costa Rica | Anonymous |
| 330 | ASSCR2867-12 | *Oncideres repandator* | Costa Rica | Anonymous |
| 331 | ASSCR2876-12 | *Oncideres senilis* | Costa Rica | Anonymous |
| 332 | ASSCR2950-12 | *Atimiola guttulata* | Costa Rica | Anonymous |
| 333 | ASSCR3060-12 | *Polyrhaphis batesi* | Costa Rica | Anonymous |
| 334 | ASSCR3136-12 | *Oreodera costaricensis* | Costa Rica | Anonymous |
| 335 | ASSCR3185-12 | *Paradiscopus maculatus* | Costa Rica | Anonymous |
| 336 | ASSCR3238-12 | *Steirastoma melanogenys* | Costa Rica | Anonymous |
| 337 | ASSCR3449-12 | *Lagocheirus rogersi* | Costa Rica | Anonymous |
| 338 | ASSCR4617-12 | *Lepturges festivus* | Costa Rica | Anonymous |
| 339 | ASSCR4716-12 | *Nyssodrysina polyspila* | Costa Rica | Anonymous |
| 340 | ASSCR4899-12 | *Colobothea aleata* | Costa Rica | Anonymous |
| 341 | ASSCR4910-12 | *Colobothea chemsaki* | Costa Rica | Anonymous |
| 342 | ASSCR4927-12 | *Colobothea distincta* | Costa Rica | Anonymous |
| 343 | ASSCR4935-12 | *Colobothea rincona* | Costa Rica | Anonymous |
| 344 | ASSCR5786-12 | *Tyrinthia moroiuba* | Costa Rica | Anonymous |
| 345 | ASSCR5814-12 | *Mimolaia calopterona* | Costa Rica | Anonymous |
| 346 | BBCCA177-12 | *Ecyrus dasycerus* | USA | Anonymous |
| 347 | BBCCM673-10 | *Lepturges confluens* | Canada | Anonymous |
| 348 | BBCCN018-10 | *Oberea perspicillata* | Canada | Hebert et al. (2016) |
| 349 | BBCCN308-10 | *Pogonocherus parvulus* | Canada | Hebert et al. (2016) |
| 350 | COLFA559-12 | *Monochamus sutor* | Finland | Pentinsaari et al. (2014) |
| 351 | COLFA576-12 | *Lamia textor* | Estonia | Pentinsaari et al. (2014) |
| 352 | COLFE1500-13 | *Tetrops starkii* | Sweden | Pentinsaari et al. (2014) |
| 353 | FBCOA919-10 | *Phytoecia nigricornis* | Germany | Anonymous |
| 354 | FBCOC126-10 | *Phytoecia icterica* | Germany | Anonymous |
| 355 | FBCOO564-13 | *Anaesthetis testacea* | Germany | Anonymous |
| 356 | JSCOL235-11 | *Leptostylus transversus* | Canada | Hebert et al. (2016) |
| 357 | JSJUL2203-11 | *Eupogonius pauper* | Canada | Hebert et al. (2016) |
| 358 | MACOL2281-12 | *Batocera granulipennis* | Pakistan | Anonymous |
| 359 | CERLF444-08 | *Astylopsis sexguttata* | Canada | Anonymous |
| 360 | CERLF453-08 | *Dectes sayi* | Canada | Anonymous |
| 361 | CERLF554-08 | *Saperda lateralis* | Canada | Anonymous |
| 362 | CERLF586-08 | *Urgleptes signatus* | Canada | Anonymous |
| 363 | GBCOU1043-13 | *Oberea pupillata* | Germany | Hendrich et al. (2015) |
| 364 | GBCOU1138-13 | *Iberodorcadion fuliginator* | Germany | Hendrich et al. (2015) |
| 365 | PSFOR986-14 | *Pogonocherus hispidulus* | France | Rougerie et al. (2015) |
| 366 | VVGPL2946-15 | *Acanthocinus sachalinensis* | Russia | Grebennikov et al. (2017) |
| 367 | COLAT119-08 | *Saperda candida* | Canada | Anonymous |
| 368 | GBMIN39969-13 | *Anoplophora horsfieldi* | China | Yang (Unpublished) |
| 369 | GBCL10096-12 | *Batocera lineolata* | Taiwan | Yeh et al. (Unpublished) |
| 370 | CER127-13 | *Atrypanius implexus* | Mexico | Anonymous |
| 371 | CER177-13 | *Lepturges angulatus* | Mexico | Anonymous |
| 372 | GBCL10088-12 | *Rondibilis horienis horienis* | Taiwan | Yeh et al. (Unpublished) |
| 373 | GBCL10092-12 | *Euseboides matsudai matsudai* | Taiwan | Yeh et al. (Unpublished) |
| 374 | GBCL10104-12 | *Anoplophora horsfieldi tonkinensis* | Taiwan | Yeh et al. (Unpublished) |
| 375 | PSFOR1134-17 | *Pogonocherus perroudi* | France | Rougerie et al. (2015) |
| 376 | ASSCR2646-12 | *Neoptychodes trilineatus* | Costa Rica | Anonymous |
| 377 | ASSCR2966-12 | *Desmiphora canescens* | Costa Rica | Anonymous |
| 378 | GBMIN48568-17 | *Lepturges angulatus* | USA | Anonymous |
| 379 | ASSCR2713-12 | *Tapeina transversifrons* | Costa Rica | Anonymous |
| 380 | ASSCR2730-12 | *Adetus costicollisAS3* | Costa Rica | Anonymous |
| 381 | ASSCR2761-12 | *Rosalba obliqua* | Costa Rica | Anonymous |
| 382 | ASSCR2792-12 | *Hesychotypa cedestes* | Costa Rica | Anonymous |
| 383 | ASSCR2834-12 | *Lochmaeocles sparsus* | Costa Rica | Anonymous |
| 384 | ASSCR2850-12 | *Oncideres fisheri* | Costa Rica | Anonymous |
| 385 | ASSCR2864-12 | *Oncideres putator* | Costa Rica | Anonymous |
| 386 | ASSCR2881-12 | *Sternycha approximata* | Costa Rica | Anonymous |
| 387 | ASSCR2952-12 | *Blabia costaricensis* | Costa Rica | Anonymous |
| 388 | ASSCR2972-12 | *Desmiphora cirrosa* | Costa Rica | Anonymous |
| 389 | ASSCR3113-12 | *Myoxinus pictus* | Costa Rica | Anonymous |
| 390 | ASSCR3150-12 | *Oreodera glauca* | Costa Rica | Anonymous |
| 391 | ASSCR3212-12 | *Pseudaethomerus maximus* | Costa Rica | Anonymous |
| 392 | ASSCR3260-12 | *Alcathousiella polyrhaphoides* | Costa Rica | Anonymous |
| 393 | ASSCR3295-12 | *Antecrurisa apicalis* | Costa Rica | Anonymous |
| 394 | ASSCR3441-12 | *Lagocheirus plantaris* | Costa Rica | Anonymous |
| 395 | ASSCR4594-12 | *Leptostylus trigonus* | Costa Rica | Anonymous |
| 396 | ASSCR4666-12 | *Nealcidion brachiale* | Costa Rica | Anonymous |
| 397 | ASSCR4687-12 | *Neoeutrypanus decorus* | Costa Rica | Anonymous |
| 398 | ASSCR4730-12 | *Nyssodrysternum serpentinum* | Costa Rica | Anonymous |
| 399 | ASSCR4764-12 | *Olenosus serrimanus* | Costa Rica | Anonymous |
| 400 | ASSCR4789-12 | *Pentheochaetes turbidus* | Costa Rica | Anonymous |
| 401 | ASSCR4814-12 | *Stenolis laetifica* | Costa Rica | Anonymous |
| 402 | ASSCR4946-12 | *Colobothina perplexa* | Costa Rica | Anonymous |
| 403 | ASSCR4975-12 | *Phaea flavovittata* | Costa Rica | Anonymous |
| 404 | ASSCR5809-12 | *Bactriola vittulata* | Costa Rica | Anonymous |
| 405 | ASSCR5811-12 | *Asemolea minuta* | Costa Rica | Anonymous |
| 406 | BBCCA3340-12 | *Astylopsis sexguttata* | USA | Anonymous |
| 407 | BBCCM001-10 | *Oberea quadricallosa* | Canada | Anonymous |
| 408 | BBCCN080-10 | *Saperda calcarata* | Canada | Hebert et al. (2016) |
| 409 | CER016-12 | *Tapeina transversifrons* | Mexico | Anonymous |
| 410 | COLFA603-12 | *Exocentrus lusitanus* | Estonia | Pentinsaari et al. (2014) |
| 411 | COLFB903-12 | *Saperda carcharias* | Finland | Pentinsaari et al. (2014) |
| 412 | COLFE001-12 | *Acanthocinus griseus* | Finland | Pentinsaari et al. (2014) |
| 413 | COLFG022-13 | *Oplosia cinerea* | Finland | Pentinsaari et al. (2014) |
| 414 | JSCOL407-11 | *Urgleptes querci* | Canada | Hebert et al. (2016) |
| 415 | PSFOR030-13 | *Acanthocinus reticulatus* | France | Rougerie et al. (2015) |
| 416 | PSFOR131-13 | *Monochamus galloprovincialis* | France | Rougerie et al. (2015) |
| 417 | CERGL086-08 | *Anoplophora glabripennis* | Canada | Anonymous |
| 418 | CERLF558-08 | *Saperda puncticollis* | Canada | Anonymous |
| 419 | COLFG061-13 | *Leiopus nebulosus* | Finland | Pentinsaari et al. (2014) |
| 420 | CERPA246-08 | *Monochamus obtusus* | Canada | Anonymous |
| 421 | GBMIN39966-13 | *Anoplophora glabripennis* | USA | An et al. (Unpublished) |
| 422 | HMCOC601-09 | *Monochamus scutellatus* | Canada | Hebert et al. (2016) |
| 423 | CER106-13 | *Acrocinus longimanus* | Mexico | Anonymous |
| 424 | GCOL12336-16 | *Exocentrus punctipennis* | Germany | Rulik et al. (2017) |
| 425 | CER124-13 | *Ecyrus lineicollis* | Mexico | Anonymous |
| 426 | GBMNE15828-21 | *Phryneta leprosa* | Malta | Mifsud and Vella (Unpublished) |
